# Single-cell and *in situ* spatial analyses reveal the diversity of newly born hematopoietic stem cells and of their niches

**DOI:** 10.1101/2024.10.14.618250

**Authors:** Léa Torcq, Catherine Vivier, Sandrine Schmutz, Yann Loe-Mie, Anne A. Schmidt

**Affiliations:** Institut Pasteur, Université Paris Cité, CNRS UMR3738, Department of Developmental and Stem Cell Biology, F-75015 Paris, France; Sorbonne Université, Collège doctoral, F-75005 Paris, France; Institut Pasteur, Université Paris Cité, Cytometry and Biomarkers UTechS/Cytometry Platform, F-75015 Paris, France; Institut Pasteur, Université Paris Cité, Bioinformatics and Biostatistics Hub, F-75015 Paris, France

**Keywords:** hematopoietic stem cells, niches, single-cell transcriptomics, MARS-seq, RNAscope, zebrafish

## Abstract

Hematopoietic stem cells (HSCs) and more committed progenitors (collectively referred to as HSPCs) emerge from vessels during development, via Endothelial-to-Hematopoietic Transition (EHT). Recently, using the zebrafish embryo, we showed that two EHT cell types emerge from the aorta, raising the question of their subsequent fate. To address this issue, we established a complex pipeline based on single-cell photoconversion and transgenic lines to characterize the transcriptomic profiles of single EHT cell type progenies. We obtained, at unprecedented resolution in the early larva, a cartography of HSPCs and highly diversified differentiated populations, notably NK-like cell types, innate lymphoid cells and early eosinophils. We show that the two EHT cell types previously characterized indeed lead to differentially fated cells, with significant differences in thymus colonization and T-lymphoid lineage commitment. Using HSPC signatures retrieved from our datasets – namely *gata2b* and *cd34/podocalyxin -,* and to address niches, we performed *in situ* gene expression analyses via RNAscope. Unexpectedly, we unveil a niche contacting the supra-intestinal artery. Finally, integration with previous datasets reveal that our populations contain potential developmental HSCs bearing signatures highly similar with adult HSCs.

**Summary Statement:** Single cell photoconversion of emerging hematopoietic precursor cells and transcriptomics unravel the diversity of hematopoietic stem and progenitor cell populations and homing in developmental niches *in toto*, during zebrafish development.

## Introduction

In vertebrates, Hematopoietic Stem Cells (HSCs), endowed with both long-term self-renewal and differentiation potential are generated during a short time-window of the embryonic development, within the intra-embryonic region named the Aorta-Gonad-Mesonephros (AGM) (Dieterlen-Lievre and Martin, 1981; Godin et al., 1993; Medvinsky et al., 1993; Thompson et al., 1998). HSC precursors (pre-HSCs) generated in the AGM emerge from a sub-population of arterial endothelium called the Hemogenic Endothelium (HE), as was suggested by studies in avian and mice (de Bruijn et al., 2002; Jaffredo et al., 1998), confirmed by *in vitro* and *ex vivo* assays (Boisset et al., 2010; Eilken et al., 2009) and visualized *in vivo* using the zebrafish model (Bertrand et al., 2010; Kissa and Herbomel, 2010). The genetic blueprint involved in the generation of HSCs is extremely conserved across vertebrates (Orkin and Zon, 2008; Yvernogeau et al., 2020). Pre-HSCs extrude from the HE according to a specific cellular process termed the Endothelial-to-Hematopoietic Transition (EHT), that has been since characterized in more details in mouse (Bos et al., 2015), chick (Sato et al., 2023) and zebrafish (Lancino et al., 2018; Lundin et al., 2020; Torcq et al., 2024). After emergence, newly-born HSC precursors are still immature, lacking repopulating ability (Batsivari et al., 2017; Rybtsov et al., 2011; Rybtsov et al., 2016). They reside in successive niches, that play a key role in driving the commitment, maturation, expansion or quiescence of hematopoietic populations. In mammals, pre-HSCs first mature in intra-aortic clusters along the ventral wall of the aorta (Taoudi and Medvinsky, 2007), before migrating to the fetal liver and finally the bone marrow where they will reside throughout life (Mikkola and Orkin, 2006). In teleosts, pre-HSCs follow a similar path, through two organs, the Caudal Hematopoietic Tissue (CHT) (Murayama et al., 2006; Tamplin et al., 2015) and the kidney marrow (Agarwala et al., 2022; Murayama et al., 2006; Traver et al., 2003), that respectively represent the functional equivalents to the fetal liver and the bone marrow of mammals.

In recent years, multiple reports highlight the heterogeneity of the HE [reviewed in (Barone et al., 2022)], that can generate both HSCs as well as multipotent or committed progenitors with more or less restricted lineage differentiation and self-renewal abilities. Progenitors emerging from extra-embryonic HE, such as erythro-myeloid progenitors (EMPs) (Chen et al., 2011; Frame et al., 2016) or T-restricted progenitors (Yoshimoto et al., 2012), have a restricted lineage differentiation potential. They are typically transient, only supporting hematopoiesis during embryonic and early life, although some of them persist throughout adulthood as tissue resident immune cells (Ghosn et al., 2019; Ginhoux et al., 2010; Gomez Perdiguero et al., 2015; Yoshimoto et al., 2011). Similarly, waves of generation of lineage restricted progenitors occur in the embryo proper [of the B- and T-lymphocyte lineages (Hadland and Yoshimoto, 2018; Yoshimoto et al., 2011)], as well as of short-term progenitors (Beaudin et al., 2014; Tian et al., 2017) and of multipotent progenitors (Dignum et al., 2021). The deciphering of all components contributing to this heterogeneity, such as the ontogeny of the HE in different regions, the transcriptional program at play in different HE subsets, the timing of emergence as well as the degree of maturation of the niches colonized by pre-HSPCs from different origins, is still in its early stages.

On this line, we recently described the co-occurrence, in the zebrafish dorsal aorta, of two types of emergence (termed EHT pol- and EHT pol+) with different morphodynamics and opposite cell polarity features (Torcq et al., 2024). These two extrusion processes stem at least partially from the differential tuning —at the single-cell level and under the control of *runx1*— of apico-basal polarity during the asynchronous maturation of the HE. Importantly, their unique cellular features, in link with their opposite cell polarity status, may impact their downstream migration and/or niching abilities, and hence their fate.

To bring light to these issues, highly specific tracing approaches coupled to transcriptomic phenotyping and simultaneous investigations of all developmental and pre-definitive niches *in toto* are required. In this context, the zebrafish embryo has been shown to be an invaluable model to study developmental hematopoiesis. In recent years and thanks to the emergence of single-cell transcriptomic technologies, many advances have been made to provide a clearer picture of the zebrafish hematopoietic cellular diversity, that parallels the diversity observed in mammals, highlighting once more the relevance of this model for fundamental and applied research. These investigations focused either on early developmental stages (Farrell et al., 2018; Sur et al., 2023; Ulloa et al., 2021; Xia et al., 2021; Xue et al., 2019; Zhao et al., 2022), focusing on HE and pre-HSCs characterization, or in adults (Athanasiadis et al., 2017; Carmona et al., 2017; Hernández et al., 2018; Hu et al., 2023; Hu et al., 2024; Kobayashi et al., 2019; Macaulay et al., 2016; Moore et al., 2016; Rubin et al., 2022; Tang et al., 2017), leaving a gap in our understanding of the developmental origins and the niches seeded by many cell populations.

Here, using single-cell photoconversion coupled to MARS-seq analysis, we show that our two newly described types of EHT are ultimately differentially fated, in particular with regards to the lymphoid lineage. In parallel, we built a multimodal sc-RNA-seq dataset of developmental HSPCs, harvesting cells from multiple transgenic lines as well as from multiple maturing niches, to draw a cartography of early hematopoietic diversity at unprecedented resolution. Finally, using whole-mount high-resolution *in situ* hybridizations, we highlight the complexity of the regionalization of the AGM and pronephros niches.

## Results

### Progenies of the two EHT cell types have different propensities to colonize the thymus

To gain insight into the fate of the two EHT cell types emerging from the aortic floor of the dorsal aorta in the zebrafish embryo (Torcq et al., 2024), we set up a single cell photoconversion protocol using the *Tg(kdrl:nls-kikume)* fish line (**Fig. 1A**). Single EHT pol+ and EHT pol-cells were photoconverted from green to red fluorescence at 2dpf (48-58hpf) (**Fig. 1A, B**) and larvae were subsequently imaged by spinning disk confocal microscopy at 5dpf (which was possible owing to the long half-life of the photoconverted Kikume, see **Fig. S1** and **Methods**). Progenies of photoconverted cells were imaged in the entire larvae and more specifically in the three niches expected to be reached at 5dpf, namely the thymus, the AGM and the CHT (**Fig. 1A**). Of notice, the definitive niche - the pronephros - was not investigated systematically since single cell photoconversion led to too few cells that could be identified as homing unambiguously in this organ of complex spatial organization (see later, ***In situ* and *in toto* gene expression analyses reveal unanticipated diversity of HSPC niches**). On average, EHT pol+ and EHT pol-cells generated similar amounts of detectable progenies, with mean values of 14.71 and 12.42, respectively (**Fig. 1C**). Based on these measurements, we could conclude that, 3 days after photoconversion, EHT cells performed on average between 3 to 5 division cycles, with no significant difference between EHT pol- and EHT pol+ progenies. We found progenies of photoconverted EHT pol- and EHT pol+ cells in the AGM (more precisely in the sub-aortic veinous plexus, **Fig. 1D**), in the CHT (**Fig. 1F**) and in the thymus (**Fig. 1H**). The highest number of progenies was found in the CHT, with mean values of 6.67 and 9.29 for EHT pol- and EHT pol+ cells, respectively (**Fig. 1G**, left), in agreement with the function of this niche in HSPC expansion (Mahony et al., 2016; Murayama et al., 2006; Tamplin et al., 2015). We also found progenies in the thymus as well as in the AGM, the latter representing lowest values (with mean values of 4.33 and 3.68 in the thymus and of 1.42 and 1.61 in the AGM, for EHT pol- and EHT pol+ cells, respectively, **Fig. 1I, E**, left). In the 8 independent experiments that we have conducted, while approximately 95% of both EHT pol- and EHT pol+ cell progenies were found in the veinous plexus of the CHT (**Fig. 1G**, right), EHT pol-progenies exhibited a lower tendency to home in the AGM (**Fig. 1E**, right: mean values of 27.78% versus 43.54% for EHT pol- and EHT pol+ progenies, respectively). Importantly, the most striking and significant difference in numbers were found for the thymus for which 100% of EHT pol-progenies reached this organ in comparison to 72.92% on average only for EHT pol+ daughter cells (**Fig. 1I**, right).

**Figure 1:**
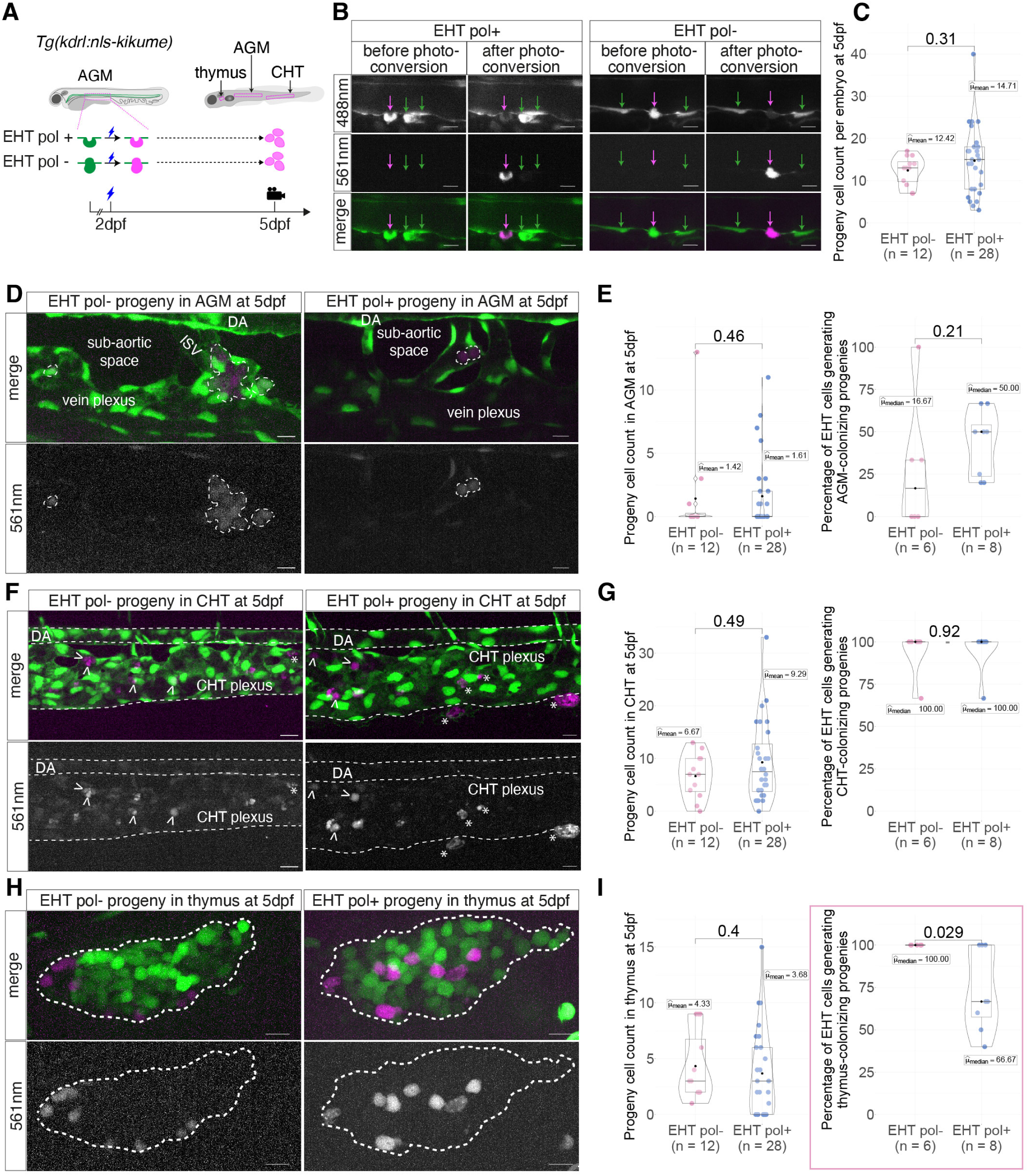
Progenies of the two EHT cell types have different propensities to colonize the thymus. **(A)** Single-cell tracing strategy to follow hematopoietic cells post-EHT, using the *Tg(kdrl:nls-kikume)* fish line. Photoconversion of single EHT pol+ or EHT pol-cells (before (green) and after (magenta) photoconversion, respectively) was performed in the AGM region. **(B)** Representative spinning disk confocal images of single EHT-cell photoconversions at 2dpf. Green and magenta arrows point at non-photoconverted and single-EHT photoconverted cells, respectively. **(C)** Total count of progenies at 5dpf and mean values. **(D, F, H)** Representative spinning disk confocal images (maximum z-projections) at 5dpf of single photoconverted EHT pol- (left) and EHT pol+ (right) progenies observed in the AGM region **(D)**, the CHT **(F)** and the thymus **(H)**. White dashed lines delineate photoconverted cells in the sub-aortic space of the AGM **(D)**, the aorta, the vein plexus **(F)** or the thymus **(H)**. White arrowheads: photoconverted cells; white asterisk: pigment cells in **(F)**. **(E, G, I)** Left plots: quantification of progenies at 5dpf in the AGM, CHT and thymus, respectively. Right plots: Percentage of photoconverted EHT cells per experiment generating AGM, CHT or thymus colonizing progenies, respectively. n=12 for EHT pol- and n=28 for EHT pol+ cells photoconverted at 2dpf, n=8 independent experiments. Two sided Wilcoxon tests. Scale bars: 10µm.

To complement our findings and in particular the significant difference in the propensity of EHT pol-versus EHT pol+ progenies to reach the thymus, we carefully analyzed the localization of these cells using 3D image analysis with the Imaris software (**Fig. 2A** and **Movie 1**). We found a significant difference in the spatial distribution of the respective progenies, with EHT pol-progeny distributed equally between central and distal regions while EHT pol+ progeny being 4.6 times more likely to be located inside the thymus than in its periphery (**Fig. 2B, C**). Finally, we reasoned that fluorescence intensity could be used as a proxy for past division events (see **Fig. S1** and **Methods**). Indeed, and theoretically, the normalized total fluorescence intensity of cells would incrementally be divided by two every cell division cycle. We thus leveraged it to gain insight into the events occurring between the photoconversion at 2dpf and implantation in the thymus at 5dpf. We did not observe any difference when globally comparing the fluorescence intensities (**Fig. 2D**), nor did we when comparing the proportion of cells relative to the number of division cycles they have been through (**Fig. 2E**).

**Figure 2:**
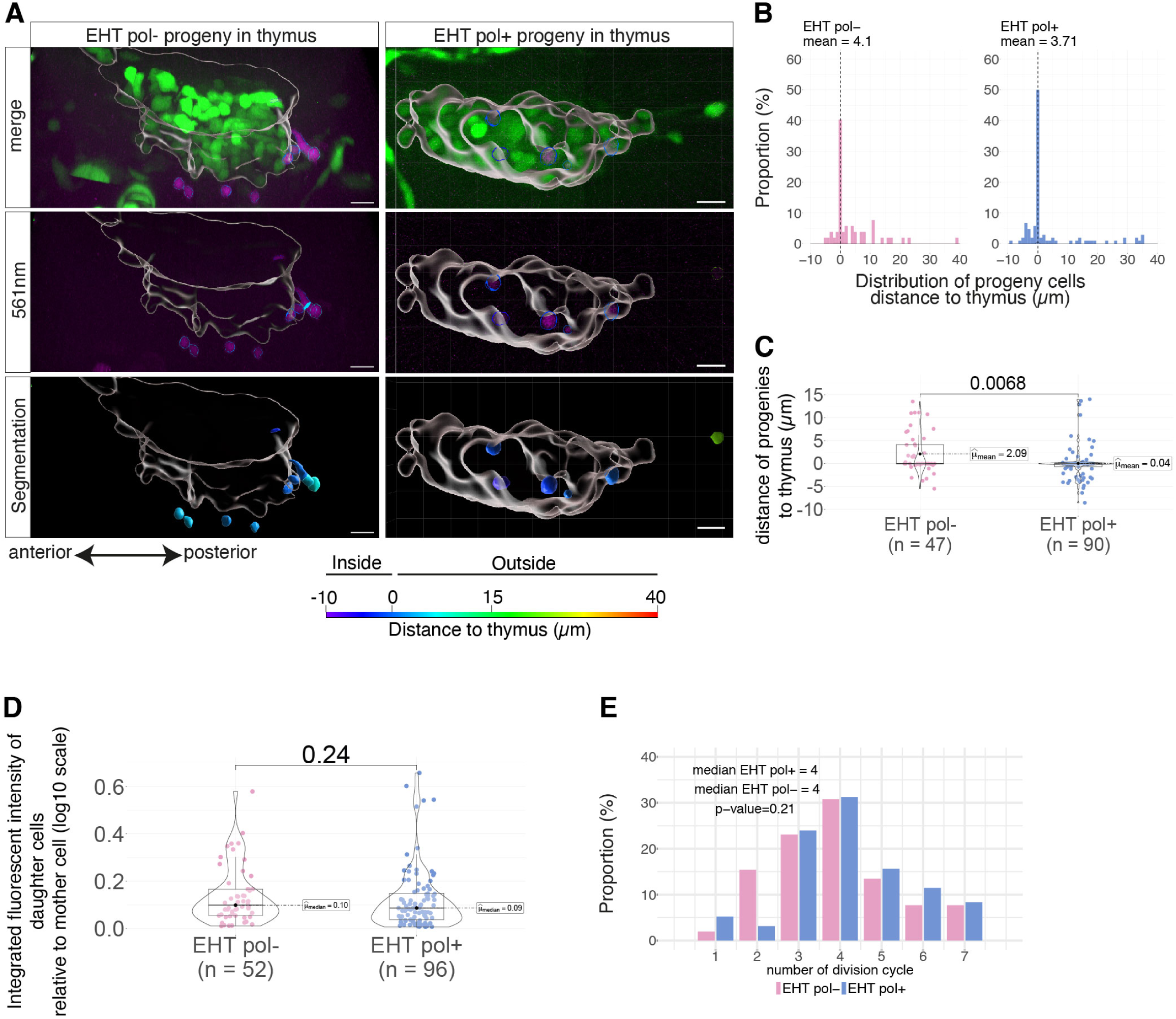
Progenies of the two EHT cell types establish in distinct thymic regions. **(A)** 3D representative surface-rendering images of thymi and progenies of single photoconverted EHT pol- (left) and EHT pol+ (right) cells observed in the thymus region of *Tg(kdrl:nls-kikume)* larvae, at 5dpf. Hematopoietic cells in the thymus (in green, cells expressing the Kikume; in magenta, the photoconverted cell progenies) were segmented and used to position the external limit of the organ. Position of progenies was measured according to a thymus distance-based color scale (scale at the bottom of the images). Scale bars: 10µm. **(B)** Distribution of minimal distances between progenies of photoconverted cells and the thymus (n=12 and n=19 thymi for EHT pol- and EHT pol+ cells, respectively). **(C)** Distribution of minimal distances between progenies of photoconverted cells and the thymus. n=47 and n=90 for EHT pol- and EHT pol+ cells, respectively (cells farther than 15µm were removed from the comparison). **(D)** Quantification of integrated fluorescence intensity (at 561nm) of photoconverted cell progenies relative to the EHT mother cell intensity, in the thymus region. **(E)** Distribution of cell division history of photoconverted cell progenies in the thymus, based on normalized fluorescence intensity at 561nm, as in **(D)**. **(B, D, E)** n=52 and n=96 cells for EHT pol- and EHT pol+ cells, respectively, n=8 independent experiments. Two sided Wilcoxon tests.

Overall, the discrepancy in the ability of EHT pol- and EHT pol+ cells to generate progenies establishing in the thymus at 5dpf as well as the uneven patterns of localization within, and in the close vicinity of, this hematopoietic organ suggest a potential impact of their cell biological features on their respective differentiation into specific lymphoid lineages (see **Discussion**).

### Single cell transcriptomics of newly born hematopoietic stem and progenitor cells reveals unexpected diversity

To gain insight into the differentiation potential of the 2 EHT cell type progenies, we performed a single cell analysis at 5dpf, after single cell photoconversion followed by MARS-seq analysis (see later). In parallel, to increase cell number and obtain a robust reference dataset at the same stage, we performed a 10XGenomics Chromium sc-RNA-seq analysis (**Fig. 3**). To enrich our dataset in hematopoietic populations and mitigate a potential reporter bias, we outcrossed two established hematopoietic reporter lines, namely *Tg(cd41:eGFP)* and *Tg(gata2b:RFP)* (**Fig. S2**). Double and single positive cells were FACS-sorted after dissection of anterior and posterior regions of 5dpf larvae that encompass, on the one side, the AGM, the thymus and the pronephros and, on the other side, the CHT (in the tail) (**Fig. 3A**). We generated 2 replicate datasets, tallying up to 11 505 analyzed hematopoietic cells for the anterior region and 14 218 for the tail. Data analysis was performed using the Seurat pipeline (Hao et al., 2021; Satija et al., 2015), and a specific integration step using rliger (Welch et al., 2019) was necessary to integrate the data from anterior and tail samples. Unsupervised clustering distinguished 42 clusters that were manually analyzed and assigned 14 distinct hematopoietic identities (**Fig. 3B**) based on the expression of specific markers (**Fig. 3C**, **Tables S1** for DEG and **S2** for references). We identified both un-committed cells, including a population of eHSPCs (embryonic/early Hematopoietic Stem and Progenitor Cells) and MPPs (MultiPotent Progenitors), as well as more selectively committed progenitors of the lymphoid, myeloid (including a population of eosinophil progenitors) and erythro-megakaryocytic lineages. Finally, we identified terminally differentiated cells, such as erythrocytes, neutrophiles, macrophages or NK-like (Natural Killer) and ILC-like (Innate Lymphoid Cells) cells. eHSPC and MPP clusters are characterized by the expression of *cmyb*, as well as either the transcription factor *gata2b*, a *cd34/podocalyxin* ortholog (*si:dkey-262h17.1*) and the DNA-binding structural protein *si:ch211-161c3.6*, a potential ortholog of HMGI-C (see below for more detailed description of our eHSPC population). Among the myeloid lineage, we could unambiguously discriminate between macrophages (expressing *spi1a/b*, *mfap4* (*mfap4.1*), *mpeg1.1*), dendritic cells (expressing *spock3* and *cd74a/b*), neutrophiles (expressing *mpx*, *cpa5* and *lyz*), and a mix of eosinophils and progenitors. The latter appear to express relatively strongly the *xbp1* transcription factor (appearing in the top 10 of DEGs, see **Table S1**), several genes of the *mfap4* locus, including an isoform of *mfap4* distinct from the one expressed in macrophages [*MFAP4 (1 of many)*, whose peptide sequence and gene identity are not precisely referenced in the Zfin database (https://zfin.org/)], as well as the lineage-specific differentiation marker *eslec* [eosinophil-specific lectin, *si:dkey-75b4.10*, that was identified recently (Li et al., 2024)]; *eslec* was found to be expressed in a very minor fraction of our eosinophils that should be the most advanced in their differentiation (**Fig. 3C**). Among the erythro-megakaryocytic lineage (expressing *gata1a*), we identified erythrocytes and erythroid progenitors (expressing *znfl2a*, *epor*, *blrvb*, *alas2* or *hbbe2*), as well as thrombocytes (expressing *mpl* and *itga2b*). Among the lymphoid lineage, we characterized T-lymphocyte progenitors (TLPs, expressing *bcl11ba*, *lck*, *runx3*, *ccr9b*, with both *rag1*+ and *rag1*-populations; for further characterization of our TLPs and comparison with juvenile/adult populations, see beneath), NK-like cells (expressing *eomesa*, *id2a*) and two ILC-like populations, populations both strongly expressing *id3*, and more specifically expressing *gata3*, *il13* and *sox13* for ILC2-like cells and *traf3ip2l* and *il7r* for ILC3-like cells **(Fig. 3C)**. Although lymphoid lineage populations have been described by sc-transcriptomic analysis in the adult zebrafish (Carmona et al., 2017; Hernández et al., 2018; Hu et al., 2023; Moore et al., 2016; Rubin et al., 2022; Tang et al., 2017), such diversity was never accessed during development (Farrell et al., 2018; Sur et al., 2023; Ulloa et al., 2021; Xia et al., 2021; Xue et al., 2019). Finally, as expected at 5dpf, we could not identify any B-lymphocyte precursors (Page et al., 2013).

**Figure 3:**
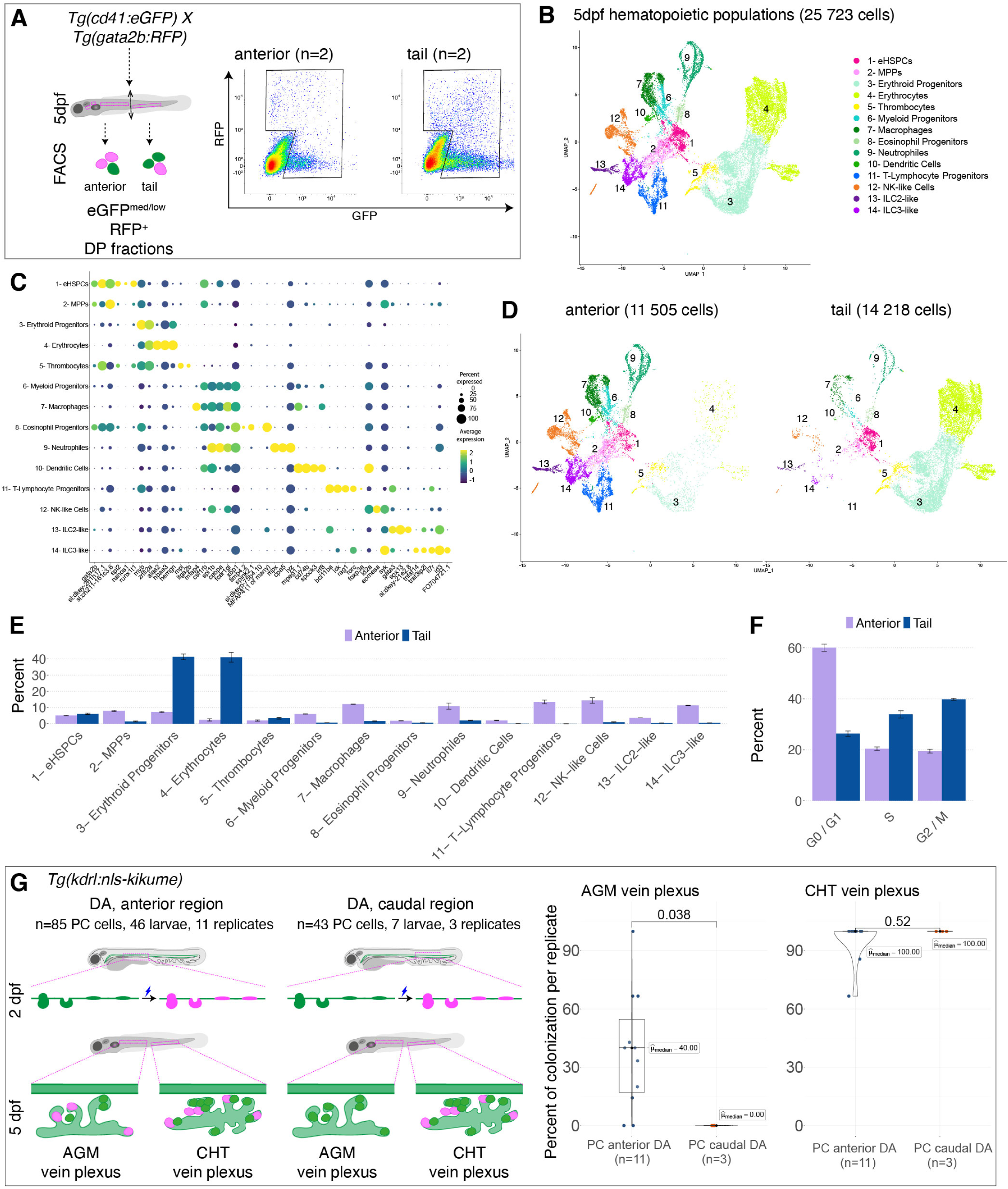
Transcriptional characterization of hematopoietic populations in the zebrafish larvae *in toto*. **(A)** Cell sorting strategy for single cell RNAseq analysis of hematopoietic populations from *Tg(cd41:eGFP; gata2b:RFP)* 5dpf larvae. A mix of single positive and double positive (DP) cells (selection gate in black on FACS plots) from either the anterior or the tail region of larvae were collected separately for Chromium assay (n=2 independent replicates per region). **(B)** UMAP visualization of 5dpf hematopoietic populations (1-14 identified clusters). **(C)** Dot-plot of selected marker genes differentially expressed across hematopoietic populations. **(D)** UMAP split view of **(B)** showing cells collected either from the anterior (left) or tail (right) regions. **(E, F)** Proportion of cell types **(E)** and cells at different cell cycle stages (**F**) in the anterior and tail regions, mean ± sd. **(G)** Experimental strategy to investigate the origins of hematopoietic cells niching in the AGM region. Using the *Tg(kdrl:nls-kikume)* fish line, HE cells and EHT-undergoing cells from the floor of the dorsal aorta were photoconverted (PC) between 35hpf and 55hpf, either in the anterior or the caudal region (left cartoons, n=85 and n=43 cells were photoconverted in the anterior and the caudal regions of n=46 and n=7 embryos, over n=11 and n=3 independent experimental replicates, respectively). Progeny in the vein plexus of the AGM and of the CHT was evaluated at 5dpf. Plots show the percentage of larvae per replicate in which progeny cells were found niching in the AGM (left) and the CHT (right) (after photoconversion of either anterior (blue) or caudal (orange) aortic floor cells). Two sided Wilcoxon tests.

To characterize further our lymphoid populations and compare with more advanced stages of differentiation, we retrieved sc-RNA-seq data from juvenile and adult thymic populations (Rubin et al., 2022) and integrated it to our own lymphoid populations, including TLPs as well as the closely related NK-like, ILC-like and dendritic cell populations (**Fig. S3A, B, and Methods**). Regarding TLPs, although our embryonic populations clustered with ETPs, maturing or cycling T-lymphocytes (**Fig. S3B**), this analysis is limited by the absence of detection, in our dataset, of T-cell surface markers, including *tcr α/β/γ/δ* or *cd8/cd3*. This suggests that our populations are early progenitors rather than maturing or differentiated populations. We characterized 3 sub-clusters, with different degrees of early differentiation and of cell cycle status (**Fig. S3C, D**). The TLP1 cluster can be considered the most upstream in the differentiation process: key transcription factors such as *gata3*, *runx3*, *bcl11ba* or *tcf7* are expressed, but *rag1* is not. The TLP2 cluster is probably more advanced in the maturation process, because of its higher expression of *cd247* [*tcrζ*, involved in early maturation steps (Levelt et al., 1993; Yoder et al., 2007)], the detection of *rag1* as well as of *cd4-1* expression. Finally, the cycling TLP clusters is highly similar to the TLP2 clusters except for its high expression of cell division related genes. In a second phase we extended our comparative analysis to our innate lymphoid populations. Our NK-like cell population appears to be more diversified than the NK population described by Rubin et al (Rubin et al., 2022) (**Fig. S3B**). We characterized 5 sub-clusters, expressing diverse immune response related genes (including granzymes, nk-lysines, chemokine-ligands or perforins, **Fig. S3E**). The thymic NK-cells described by Rubin et al. seem to correspond in majority to our sub-clusters 1 and 5, suggesting our NK sub-cluster heterogeneity might reflect the niche diversity we access with our *in toto* approach. Indeed, markers of our sub-clusters 3 and 4 (e.g. *ccl38.6, gzm3, vav3b*), although not detected in thymic NK-cells of the Rubin et al dataset, are found to be expressed in other NK-cells harvested from diverse tissues described in the literature, such as spleen NK-cells (Hu et al., 2023), kidney marrow NK-cells (Tang et al., 2017) or whole-body NK-cells (Carmona et al., 2017; Hernández et al., 2018). Finally, we characterize two terminally differentiated ILC clusters (**Fig. S3C, F**), ILC2-like and ILC3-like, demonstrating for the first time their presence at such an early stage. They express specific type 2 and type 3 immune response interleukins (respectively *il13/4* for ILC2-like and *il22/17* for ILC3-like cells) as well as lineage specific transcription factors (respectively *gata3* and *rorc*) and ILC-lineage differentiation inducing transcription factor *id2a*.

Overall, using transcriptomic assay and known markers of each population (see **Table S2** for reference for key markers), we describe for the first time the unexpected diversity of zebrafish early larval hematopoietic populations.

Comparing the UMAPs obtained with the two regionally restricted datasets (**Fig. 3D**), we observed a clear difference in the representation of hematopoietic populations in the anterior and tail regions. Quantification shows that the tail contains a large majority of erythroid cells (85.6% ± 5.2, with equal amounts of progenitors and differentiated cells), as well as eHSPCs/MPPs (7.5% ± 0.7), with very little representation of cells of the myeloid (5.0% ± 0.5) and lymphoid (1.9% ± 0.3) lineages (**Fig. 3E**). In comparison, the anterior region harbors a wider diversity of cell types, in more homogeneous proportions, with 13.0% ± 0.5 of eHSPCs/MPPs, 11.6% ± 1.4 of erythroid cells, 32.7% ± 2.3 of myeloid cells and 42.7% ± 2.9 of lymphoid cells (**Fig. 3D, E**). Gene-based analysis of cell cycle-state revealed that 3/4 of the cells in the tail were actively cycling (with 33.8% ± 1.4 in S phase and 39.8% ± 0.4 in G2/M phase, **Fig. 3F**), in agreement with its role in supporting the expansion of newly generated hematopoietic cells (Murayama et al., 2006). Conversely, cells in the anterior region are in majority arrested in the cell cycle (G0/G1, 60.1% ± 1.4, **Fig. 3F**). This suggests that the AGM and CHT environments provide different signaling cues for quiescence versus division, in particular for erythroid populations that represent almost 90% of cycling cells in the tail region (see **Fig. S4A**, and **Source data Table**). Importantly, it also reinforces the idea that the CHT is a significant niche for eHSPCs (representing 5.1 and 6.1% of the anterior and tail cells respectively), in agreement with the previously described functional relationship between HSPCs and the vein plexus (Tamplin et al., 2015).

### The AGM provides a specific niche for HSPCs newly born from the anterior part of the dorsal aorta

The relatively high diversity of cell populations in the anterior region, together with previous studies showing that the dorsal aorta in the trunk region provides long-term hematopoietic populations [which is not the case for the cells emerging from the aorta in the tail region, see (Jin et al., 2009; Tian et al., 2017)], raised the possibility of the AGM as a niche providing specific cues, different from the CHT. Also, it was determined that cells emerging in the AGM region reach the CHT after migration throughout the vein and transport via circulation (Jin et al., 2007; Murayama et al., 2006); however, the origin of cells niching in the AGM vascular plexus appears to be more elusive, with supposedly some cells implanting in the sub-aortic space after having passed by the CHT. To bring further insight into these issues, we photoconverted hemogenic cells in the floor of the dorsal aorta either in the trunk region, or in the tail (the caudal artery), at 2dpf (**Fig. 3G**), and assessed the presence/absence of progenies both in the AGM and the CHT vein plexus at 5dpf. In both cases, progeny cells were observed in the CHT at 5dpf, with no difference in frequency (medians of 100%, **Fig. 3G** right plot). However, we did not observe any progeny of cells emerging from the aorta in the CHT colonizing the AGM region (**Fig. 3G**, left plot) while we visualized an average of 38% of the progeny of cells emerging in the AGM homing in this region, 3 days after photoconversion (**Fig. 3G**, left plot). Furthermore and importantly, in the case of cells photoconverted in the dorsal aorta in the trunk region, we observed progenies homing in the direct vicinity of the region of emergence (see **Methods**). Altogether, these data are in favour of the idea that the AGM is a primary niche into which a significant proportion of newly born HSPCs home without necessarily passing by the CHT (see also **Discussion**).

### The progenies of EHT pol+ and EHT pol-cells are multipotent but are endowed with different lymphoid differentiation potential

Differences in the ability of EHT pol+ and EHT pol-progenies to colonize the thymus prompted us to characterize their identity at the transcriptomic level. For this, we performed single-cell transcriptomics on index-sorted progenies of single-photoconverted EHT cells (**Fig. 4A**, condition 1), using the MARS-seq protocol (Jaitin et al., 2014; Keren-Shaul et al., 2019). In addition to progenies of single photoconverted EHT cells, and to mitigate the low number of cells such a highly specific approach provides, we combined additional modalities in our MARS-seq dataset (**Fig. 4A**, **Table S3**). To increase the number of progenies of cells photoconverted in the aortic floor while undergoing the EHT, we performed photoconversion of the HE (**Fig. 4A**, condition 2). To add spatial information, we also dissected the 5dpf larvae (3 days after photoconversion and prior to cell dissociation) to obtain cells either from the anterior region and the tail (**Fig. 4A**, condition 2, bottom right), or from the head containing the developing thymus and pronephros, the trunk, and the tail regions (**Fig. 4A**, condition 2, bottom left). Additionally and to encompass the entire hemogenic sites of the larvae as was the case for the Chromium dataset (ex: that also include the dorsal aorta in the tail region), we collected hematopoietic cells using a *Tg(runx1+23:eGFP)* line that we have generated to optimize fluorescence signals, (**Fig. 4A**, condition 3). Finally, and to ease the comparison with our previously established Chromium dataset, we added a final condition using the *Tg(cd41:eGFP) x Tg(gata2b:RFP)* outcross (**Fig. 4A**, condition 4). In total, we analyzed 5119 cells from 28 replicates, that were merged for analysis, unsupervised clustering and manual annotation. Despite the complexity of this dataset in terms of cell and spatial origin as well as replicate amount, we observed strong similarities compared to our Chromium dataset. Based on the expression of marker genes, cell type assignation to clusters was practically identical to our Chromium dataset (**Fig. 4B, C** and **Table S4**), allowing us to characterized once again a wide diversity of hematopoietic cells. The merging of our two datasets (Chromium and MARS-seq) and the conservation of a similar architecture and clustering (**Fig. S5** and **Table S5**) validated the robustness of our classification across different technical and biological modalities.

**Figure 4:**
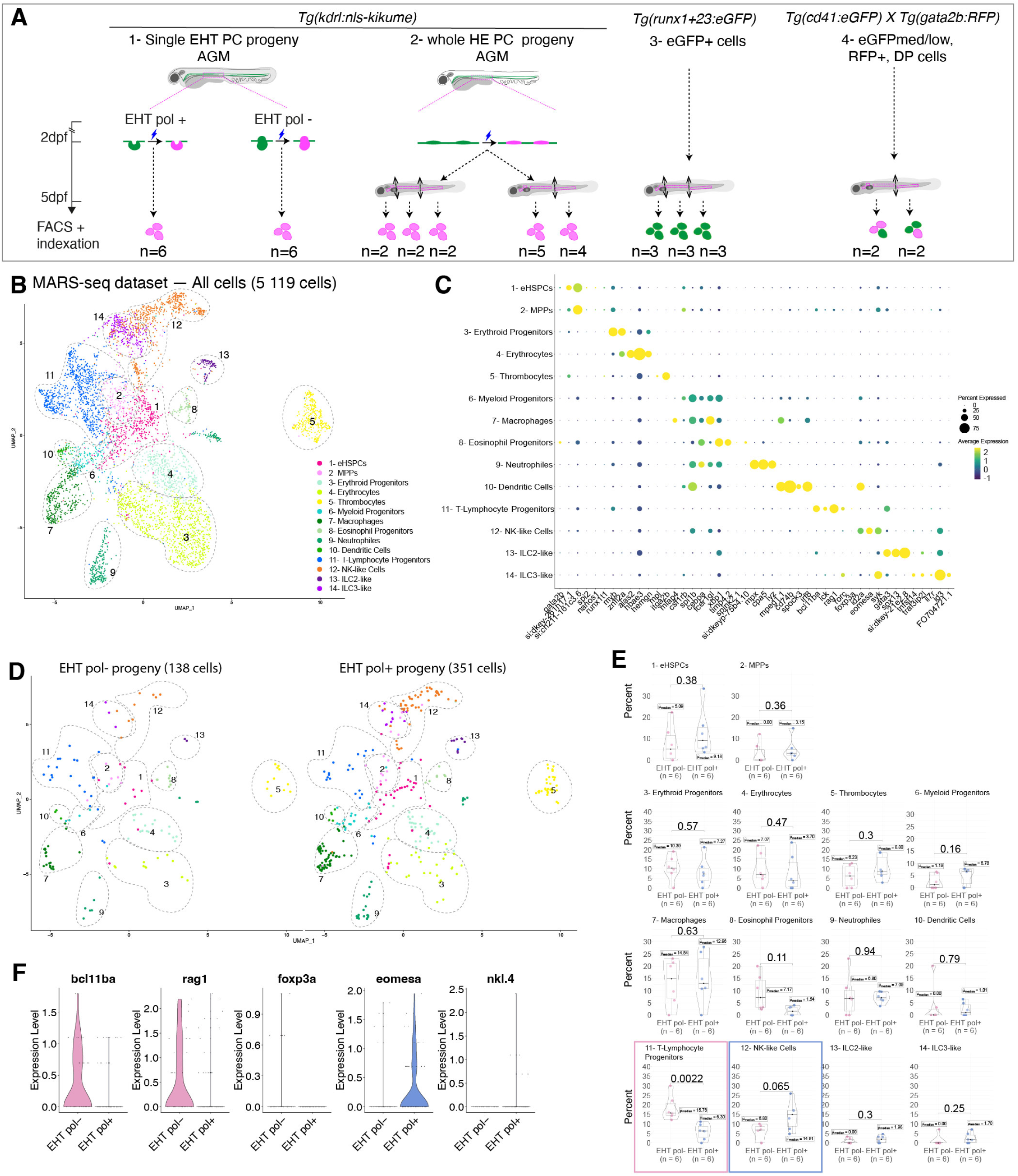
The progenies of EHT pol+ and EHT pol-cells are multipotent but are endowed with different lymphoid differentiation potential. **(A)** Schematic illustration of the single-cell collection strategy for the MARS-seq assay. Cells were collected at 5dpf according to 4 different modalities (**Table S3** for cell count per replicates). Cells from conditions 1-2 were obtained using the *Tg(kdrl:nls-kikume)* transgenic line, after performing photoconversions in the trunk region at 2dpf of either 1-EHT pol+ or EHT pol-cells, 2-HE cells, and collecting progenies at 5dpf. Progeny cells from rostral, anterior, trunk and tail regions were collected separately for condition 2. Condition 3: *Tg(runx1+23:eGFP)* GFP^+^ cells were collected from rostral, trunk and tail regions. Condition 4: *Tg(cd41:eGFP; gata2b:RFP)* eGFP^med/low^/RFP^-^, eGFP^med/low^/RFP^+^ and eGFP^-^/RFP^+^ cells were collected from anterior and tail regions. **(B, D)** UMAP visualization of hematopoietic cells from the MARS-seq dataset. **(C)** Dot-plot of selected marker genes differentially expressed across defined hematopoietic populations. **(D)** Split view from **(B)**, showing only the progenies of single-photoconverted EHT pol-cells (left) and EHT pol+ cells (right). **(E)** Proportion of each hematopoietic lineage generated by each EHT cell type, n=6 independent experiments with cells sorted on plates each time (and frozen until further use, see Methods), two sided Wilcoxon tests. **(F)** Violin plots of gene expression levels in lymphoid populations (clusters 11 to 14) within EHT pol- and EHT pol+ progenies.

Analysis of EHT pol+ and EHT pol-cell progenies revealed that both generated multipotent cells, that give rise to virtually all of the cell populations we previously identified (**Fig. 4D**). While no difference was observed in the relative proportion of most cell types (**Fig. 4E**), our data reveals a significant difference in their ability to generate cells of the lymphoid lineage (**Fig. 4E**, pink and blue boxes). While EHT pol-cells generated more than twice more T-lymphocyte progenitors than EHT pol+ cells (median values of 15.76 versus 6.30 cells, respectively), they generated more than twice less NK-like cells than EHT pol+ cells (median values of 6.80 versus 14.91 cells, respectively). This difference is apparent when comparing the expression of specific genes related to the T-lymphocyte lineage identity acquisition, such as the transcription factor bcl11ba (Bajoghli et al., 2009; Tydell et al., 2007) or the recombination regulating gene rag1 (Willett et al., 1997), that are expressed in higher proportions of EHT pol-cell progenies (**Fig. 4F**). Conversely, the transcription factor eomesa (Rubin et al., 2022), as well as the NK-specific anti-microbial peptide gene nkl.4 (Pereiro et al., 2015), are not expressed or in lower proportions by EHT pol-cell progenies (**Fig. 4F**). In line with evidence showing the existence in the embryo of multiple waves of T-cell generations (Mold et al., 2010; Tian et al., 2017), and that HSC-independent T-cells express *foxp3a* (Tian et al., 2017), a master regulator of the T-regulatory lineage, we looked for the presence of T-regulatory cell progenitors within our single-EHT progenies. Interestingly, only EHT pol-TLP progenies express *foxp3a* (**Fig. 4F**). Furthermore, the percentage of TLPs expressing *foxp3a* is higher in EHT pol-derived TLPs (22.2%) than HE or EHT derived TLPs (5.1% and 0% respectively), suggesting a higher tendency for EHT pol-to generate T-regulatory cells.

Overall, these observations are consistent with our results relating to the increased frequency of thymus colonization by progenies of EHT pol-cells (**Fig. 1I**), reinforcing the significance of the difference in the differentiation potential of the two EHT populations, in particular in regard to the lymphoid lineage.

### Developmental organs differentially support the expansion and differentiation of hematopoietic cell lineages

Quantitative analysis of the spatial distribution of cell types throughout our MARS-seq datasets (see **Fig. S6A**, cartoon) reinforced the aforementioned enrichment of erythroid cells in the tail region as well as the unexpected diversity and homogeneous representation of hematopoietic cells in the trunk region specifically (**Fig. S6A**). Particular innate lymphoid cell types such as ILC2-like, are notably enriched in the trunk region (**Fig. S6A, E, F** and **Source Data Table**). This points to the potential diversity of functional niches there (as revealed by *in situ* spatial analysis of hematopoietic cell populations using RNAscope, see beneath). In addition, undifferentiated cells such as eHSPCs and MPPs that, altogether, represent 17.5% ± 8.5 of all trunk cells, are 1.5 times more abundant in the trunk than in the tail or in the rostral region (11.8% ± 8.5 and 10.8% ± 5.9 respectively, see **Fig. S6A** and **Source Data Table**). In accordance with our Chromium dataset, we find that cells in the anterior region are less actively cycling compared to the tail (**Fig. S6B**). Here, our spatial analysis refined to the rostral and trunk regions of the anterior part of the larval body does not show any difference in the cell cycle activation between these regions, suggesting that the trunk, in contrary to the CHT, does not particularly promote the rapid expansion of newly generated hematopoietic precursors but rather contains an important proportion of non-cycling cells (similarly to the definitive hematopoietic organs in the anterior region). Comparative analysis of the hematopoietic populations identified amongst cells labelled by different transgenic lines as well as progenies of photoconverted cells (HE) reveals that identified populations are recovered by all three approaches, in relatively similar proportions (**Fig. S6C, D**). This emphasizes once more the ability of newly born hematopoietic precursor cells recovered with our photoconversion approach to generate cells of all hematopoietic lineages in a short amount of time (3 days), including progenitor cells as well as terminally differentiated ones (**Fig. S6E, F**). Finally, decomposition of our HE cells subsets into regional components (head, anterior, trunk and tail regions, **Fig. S6F**) validate that virtually all TLPs home in the most rostral region, most probably in the thymus, as opposed to other cell types that partition rather equally between the head and the trunk region (with the exception of ILC2-like cells).

Overall, together with our aforementioned results showing the local origin of cells niching in the sub-aortic region, our data suggest a specific, unforeseen role, of the AGM/trunk region in the specification and maturation of sub-sets of hematopoietic populations during zebrafish development.

### eHSPC populations contain potential HSC candidates

In both our Chromium and MARS-seq datasets, we were able to discriminate between two non-committed populations (eHSPCs and MPPs, **Fig. 5A, B**), that share similar data architecture as well as relatively well conserved gene-specific expression (**Fig. 5C**). These populations are characterized by the expression of established markers of HSCs, including *gata2b* (Butko et al., 2015; Macaulay et al., 2016), the *cd34/podocalyxin* ortholog (*si:dkey-261h17.1*) (AbuSamra et al., 2017; Rubin et al., 2022), *meis1b* (Kobayashi et al., 2010), *spi2*, *nanos1*, *fli1a* (Rubin et al., 2022) and *si:ch211-161c3.6* (the latter only being more expressed in MPPs than in eHSPCs). Of note, *si:ch211-161c3.6* was also found to be the top DEG in the HE/HSC cluster of a published early embryonic (1-2dpf) hematopoietic cell dataset (Ulloa et al., 2021). Spatial information from our MARS-seq dataset suggests that, at 5dpf, both MPPs and eHSPCs are present in all developmental niches, with cells found in the tail, the trunk and the rostral region in relatively similar proportions (**Fig. S7A**). Cell cycle analysis shows that only a fraction of eHSPCs are arrested in the cell cycle (median of 28.17%), contrary to MPPs that are in majority arrested in the cell cycle (66.67%, **Fig. S7B**). This suggests a heterogeneity within our eHSPCs cluster, with potential sub-clusters that differentially progress through the cell cycle. To complement the characterization of our developmental HPSCs, we extended our exploration of the published sc-RNA-seq datasets of Rubin et al more specifically focused on the adult zebrafish whole kidney marrow and integrated the adult and larval HSPC/MPP populations (**Fig. 5D**). Unsupervised clustering resolved 7 different clusters that, after gene expression analysis (**Table S6**), were highly similar to the clusters defined in Rubin et al (**Fig. 5E, F**). Among the 7 clusters, that all contain a mix of embryonic and adult cells (**Fig. 5F**), 3 primed populations (clusters 5-6-7 **Fig. 5E**) were characterized by the expression, at a low-level, of early lineage specific transcription factors (**Fig. S8A, Table S6,** for erythroid-, lymphoid-and myeloid-biased populations). Similarly, we characterized, in accordance with the original annotation, 2 cycling populations (clusters 2-3), with cells expressing marker genes of G2/M and S phase respectively. Finally, two additional clusters, that do not express any lineage-specific transcription factors, were characterized (clusters 1 and 4). Similarly to our dataset, one of the clusters (1, HSCs, **Fig. 5G, S8B**), shows a higher expression of key genes (such as *nanos1, meis1b*) that have been characterized in the literature as crucial for populations with long-term replenishment potential, suggesting that this cluster might represent *bona fide* HSCs. The other cluster (4, MPPs) might represent an intermediary population of multipotent progenitors. In addition to this integrative analysis using adult cells, we re-analyzed an earlier embryonic (2dpf/3.5dpf/4.5dpf) dataset (Xia et al., 2021) but failed to find transcriptionally equivalent cell populations. In particular, we were unable to recover cells co-expressing key HSCs genes, such as *gata2b*, *si:dkey-261h17.1/cd34*, *nanos1* and *spi2* (data not shown). We also looked for the expression of aforementioned HSCs signatures within progenies of single-EHT photoconverted cells (**Fig. 5H**). Although the absence of expression of 3 out of 4 markers within EHT pol-derived eHSPCs does not allow us to draw any conclusion (possibly due to a low sampling), the expression of all of those genes within progenies of EHT pol+ suggests an ability of EHT pol+ cells to generate putative *bona fide* HSCs. Overall and importantly, this shows that the 5dpf larval HSPC populations are transcriptionally closely akin to adult kidney marrow HSPCs (**Fig. S8C**), including a population of potential HSCs with the conserved *nanos1/spi2/meis1b* as well as *cd34* (*podocalyxin*) and *gata2b* signatures.

**Figure 5:**
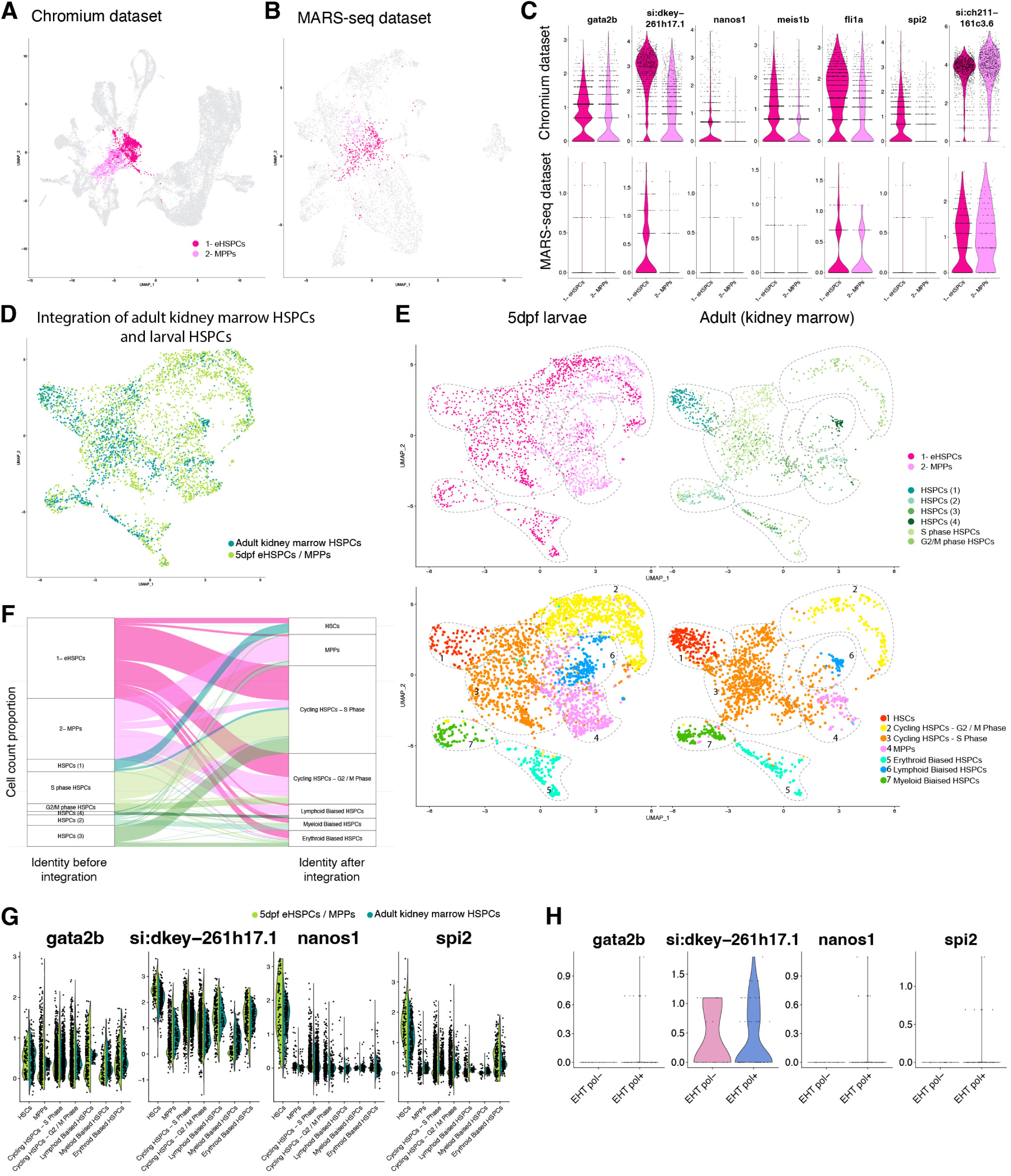
Single-cell data comparison and integration of multimodal analyses of HSPC/MPP populations. **(A-C)** Characterization of eHSPC and MPP clusters. **(A, B)** UMAPs highlighting eHSPC and MPP clusters in the complete Chromium and MARS-seq datasets respectively. **(C)** Violin plots of selected eHSPCs and MPPs marker genes in the two datasets. **(D, E)** UMAPs of integrated 5dpf whole larvae and adult kidney marrow (Rubin et al., 2022) HSPC/MPP populations, with cell populations colored by origin in **(D)**. **(E)** split view from **(D)**, with cell colored depending on origin (top, corresponding to our manual annotation for larval cells and annotations from Rubin et al. for adult cells) or cell type assignation after integration (bottom UMAPs, split view). **(F)** Alluvial plot showing the repartition of cells from larval and adult origins within integrated clusters. **(G)** Violin plots of gene expression levels of selected HSPC markers, expressed in both larval and adult datasets. **(H)** Violin plots of gene expression levels in eHSPC and MPP clusters within EHT pol- and EHT pol+ progenies.

### *In situ* and *in toto* gene expression analyses reveal unanticipated diversity of HSPC niches

The small size of the zebrafish larva offered the unique opportunity to investigate the localization of hematopoietic populations in their respective niches, in the entire body. During the developmental period (embryonic and early larval stages), the best described HSPC niches have been localized at the ultra-structural level in anterior/rostral pre-definitive hematopoietic organs such as the pronephros (Agarwala et al., 2022), as well as the CHT in the posterior/tail region (Tamplin et al., 2015).

To bring further insight into these issues, we investigated the sites of implantation of eHSPC populations in the entire body at 5dpf and performed RNAscope that allows combining high sensitivity mRNA detection with high resolution fluorescence confocal imaging.

First, we investigated the localization of hematopoietic cells expressing *gata2b*, based on the expression of this transcription factor in the eHSPC/MPP populations that we have identified in the larva at 5dpf (**Fig. 5C, G**), as well as its previously described expression in zebrafish pan-HSPC populations [in particular, *gata2b* was proposed to be one of the best markers for marrow HSPCs in the adult kidney (Kobayashi et al., 2019; Rubin et al., 2022)]. We took advantage of the relative stability of the eGFP after formaldehyde/MeOH fixation of the larvae (see **Methods**) to superpose directly the fluorescence signals amplified from the probes to either vessels/endothelial cells or hematopoietic cells using two transgenic fish lines: *Tg(kdrl:eGFP)* and our *Tg(runx1+23:eGFP)* used for the MARS-seq analyses (which faithfully labels eHSPCs, **Fig. S6D** and see also (Tamplin et al., 2015)). Cells expressing *gata2b* were found in most hematopoietic tissues except for the thymus in which we observed virtually no *gata2b*+/eGFP+ cells (data not shown, **n=4**), indicating that early thymic progenitors can be distinguished from HSPCs on this basis (which is also in agreement with (Rubin et al., 2022), in the juvenile and adult organs). The other hematopoietic organs in which we found cells expressing *gata2b* include the pronephros (**Fig. 6A**) in which they lodge relatively homogenously in regard to the territory occupied by all eGFP+ cells (**Fig. 6B**, quantifications based on 3D reconstitutions using the Imaris software; see also 3D rotations in **Movie 2,** n=3 larvae shown), the AGM and the CHT (**Fig. 6C-E**). In addition, quantitative analyses revealed that *gata2b* is expressed relatively homogenously in 24.64%, 17.19%, and 20.45% of eGFP+ cells localized in the pronephros, AGM (trunk) and CHT regions respectively (**Fig. 6F**). In the 3 regions, we observed that *gata2b*+/eGFP+ cells are relatively dispersed among surrounding cells (for the pronephros, see **Movie 2**) or among hematopoietic clusters (see for example the delineated clusters in the AGM **Fig. 6D** and **Movie 3** for the trunk and CHT regions, n=2 larvae shown).

**Figure 6:**
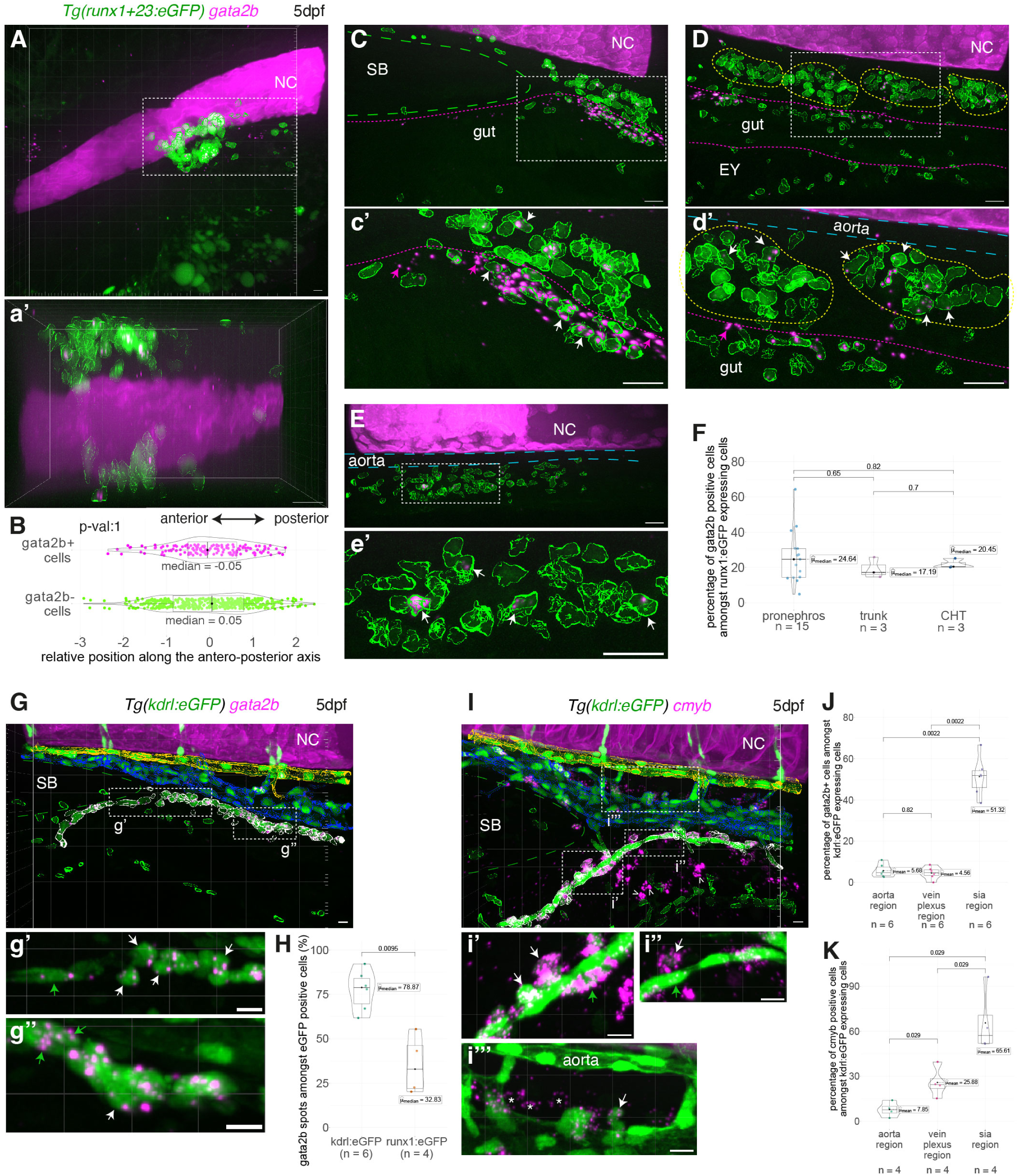
Whole mount *in situ* analysis of *gata2b* and *cmyb* mRNA expression in vascular and hematopoietic cells using RNAscope. **(A, C-E)** Representative fluorescence confocal images (Imaris 3D-rendering) of RNAscope ISH for *gata2b* (magenta spots) in 5dpf *Tg(runx1+23:eGFP)* larvae. Images show respectively the pronephros region **(A)**, the anterior trunk region (**C,** posterior to the swim-bladder), the posterior trunk region (**D,** above the elongated yolk) and the CHT **(E)**. **(a’, c’, d’, e’)** are magnifications of regions outlined with white dashed boxes in the respective panels. Hematopoietic eGFP+ cells were segmented (green contours). White arrows point at *gata2b* positive hematopoietic cells. Magenta arrows point at *gata2b* spots outside hematopoietic cells in the SIA region (see also **Movies 2-4** for resolving the information in 3D). The aorta, the swim-bladder, the gut, sub-aortic clusters, are delimited by long blue, long green, magenta and yellow dashed lines, respectively. **(B)** Relative position of eGFP+ cells along the antero-posterior axis of the pronephros (n=447 *gata2b*-cells, n=145 *gata2b*+ cells). **(F)** Percentage of hematopoietic cells expressing *gata2b*, n=15 larvae for the pronephros, n=3 for trunk and CHT regions. **(G, I)** Representative images (Imaris 3D-rendering) of RNAscope ISH for *gata2b* (**G**, magenta spots; see also **Movie 5**, top panels) and *cmyb* (**I**, magenta spots; see also **Movie 5**, bottom panels) in 5dpf *Tg(kdrl:eGFP)* larvae. Images show the anterior trunk region (posterior to the swim-bladder). **(g’, g’’, i’-i’’’)** are magnifications of regions outlined with white dashed boxes in **(G, I)** respectively. Vascular and hematopoietic eGFP+ cells were segmented and classified: yellow for aortic cells, blue for veinous cells and hematopoietic cells in the vascular plexus, white for cells in and around the SIA (see **Methods**) and green for non-classified cells. **(g’, g’’)** White and green arrows point at eGFP+/*gata2b*+ round cells (potentially hematopoietic) and vascular elongated cells, respectively; **(i’-i’’’)** white and green arrows point at eGFP+/cmyb+ hematopoietic and vascular elongated cells, respectively; white asterisks point at eGFP-/*cmyb+* potential hematopoietic cells (also visible in the gut region, white arrowheads in (I)). **(H)** Percentage of *gata2b* mRNA spots located in hematopoietic cells (using the *Tg(runx1+23:eGFP)* background, n=4) or in vascular/newly generated potential hematopoietic cells (using the *Tg(kdrl:eGFP)* background, n=6). **(J, K)** Percentage of vascular and hematopoietic eGFP+ cells expressing *gata2b* or *cmyb,* respectively (n=6 and n=4). **(B, F, H, J, K)** Two sided Wilcoxon tests. NC: notochord, SB: swim-bladder, EY: elongated yolk. Scale bars: 10µm.

Finally and quite importantly, we noticed, in the anterior part of the trunk region, beneath the vein and in proximity to the gut, a population of *gata2b*+ putative cells that do not express eGFP (**Fig. 6C**, see the magnified area in **Fig. 6c’**, with the white arrows pointing at the few *gata2b*+/eGFP+ cells surrounding the area containing *gata2b*+/eGFP-putative cells (magenta arrows); see also **Movie 4,** n=3 larvae shown, imaged in the upper AGM/trunk region). We then investigated if these *gata2b* signals may be associated with vascular structures, as they appear to align longitudinally to - and above - the gut, in the region of the Supra-Intestinal Artery [SIA, (Isogai et al., 2001)]. RNAscope performed with the *Tg(kdrl:eGFP)* fish line revealed that indeed, *gata2b* signals co-localize with eGFP+ endothelial cells of the SIA (**Fig. 6G** and magnifications **Fig. 6g’, g’’** (green arrows) and, in addition, with cells in its close vicinity (**Fig. 6g’, g’’** (white arrows); see also **Movie 5**, top panels, n=3 larvae at 5dpf, imaged in the upper AGM/trunk region). Quantitative analyses revealed a very significant increase in *gata2b* signal co-localization with kdrl:eGFP+ cells in comparison to runx1:eGFP+ cells in this region of the trunk (**Fig. 6H**, with 78.87% and 32.83% for kdrl:eGFP+ and runx1:eGFP+ cells, respectively), as well as in the percentage of cells that are double positive *gata2b*+/eGFP+ in the near vicinity or contacting the SIA, in comparison to the dorsal aorta and vein plexus regions (approximately ten times more, **Fig. 6J**). Apart from SIA endothelial cells expressing *gata2b*, some of the *gata2b*+/eGFP+ cells that surround the SIA could possibly be young MPPs (our single-cell results indicate that a minor fraction of the MPP population expresses *gata2b* at a higher level than eHSPCs, **Fig. 5G**); they could also be ILCs and/or their early progenitors (note that, also in our single-cell results, a fraction of ILC2-like cells express *gata2b*, see for example our integrated data **Fig. S5C**). RNAscope using *Tg(kdrl:eGFP)* and a *cmyb* probe confirmed that some of the cells in direct contact with the SIA are double positive *cmyb+*/eGFP+, which confirms their hematopoietic nature (**Fig. 6I** and magnifications **Fig. 6i’-i’’**; see also **Movie 5,** bottom panels, n=3 larvae shown). Finally, quantitative analyses confirm that, in comparison to the aortic (dorsal aorta) and vein plexus regions, the SIA area is significantly enriched in double positive *cmyb+*/eGFP+ cells (**Fig. 6K**, also with the vein plexus containing significantly more double positive cells than the aortic region, consistently with the involvement of the vein in niche functions).

Altogether, these data suggest that, at 5dpf, the SIA is a potential niche for hematopoietic cells originating from the vascular system (see **Discussion** for additional comments).

Lastly, and in line with the expression of *gata2b* in small sub-populations as mentioned before (MPPs and of ILC2-like cells), we also looked at eosinophil progenitors that we have identified in our sc-RNA-seq data set, at 5dpf (**Figs 3C**, **S5C**). To visualize the localization of these progenitors, we chose *timp4.2*, a metallo-protease inhibitor that is a selective marker of this granulocytic cell population (**Figs 3C**, **S5C**). In addition, in regard to niches, we noticed that this population of cells is uniquely characterized by the expression of several genes whose function is potentially related to the niche/extracellular matrix such as *MFAP4 (1 of many)* or serine peptidase inhibitors of the *spink2* family. RNAscope performed with a *timp4.2* probe and using the *Tg(runx1+23:eGFP)* fish line highlighted, in comparison to *gata2b*+, the unique feature of double positive *timp4.2+/*eGFP+ cells that accumulate in the anterior part of the pronephros, in comparison to the territory occupied by all eGFP+ cells (**Fig. S9A, B;** see also **Movie 6**). This cell population is also characterized by relatively few cells lodging among hematopoietic clusters in the trunk and the CHT (**Fig. S9C, D**), with an average of 6.98%, 5.50% and 4.31% of *timp4.2* + cells among all other eGFP+ cells in the pronephros, trunk and CHT regions, respectively (**Fig. S9E**). Altogether, these last results suggest that the pronephros niche is characterized by geographical and population-specific functional properties.

We then investigated the localization of hematopoietic cells expressing *cd34/podocalyxin* (*si:dkey-261h17.1*) that is one of the best markers to identify HSPCs and, also, to discriminate between HSCs and MPPs (it is strongly expressed in HSCs, at relatively homogenous levels, see **Figs 5G**, **S8B**). Of notice, *cd34/podocalyxin* is as well strongly expressed in other cell types (<u>see Daniocell,</u> (Sur et al., 2023), https://daniocell.nichd.nih.gov/index.html), particularly endothelial and vascular smooth muscle cells which is why the glomerulus in the pronephros region is labelled (**Fig. 7A**) as well as the aorta and inter-segmental vessels (**Fig. 7C, D**).

**Figure 7:**
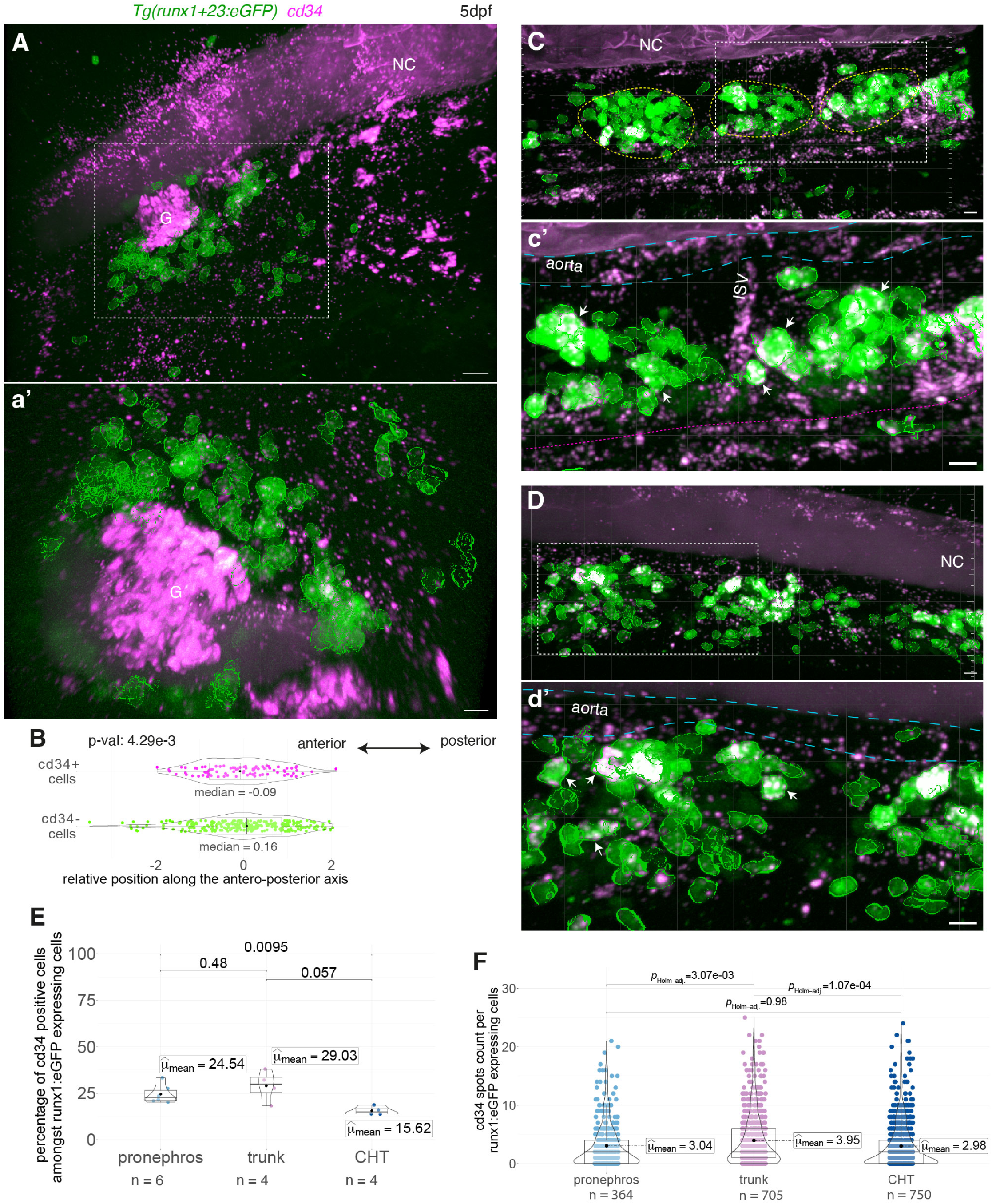
Whole mount *in situ* analysis of *cd34* mRNA expression in hematopoietic and vascular cells using RNAscope. **(A, C, D)** Representative images (Imaris 3D-rendering) of RNAscope ISH for *cd34* (magenta spots) in 5dpf *Tg(runx1+23:eGFP)* larvae. Images show the pronephros region (**A**, see also **Movie 7**), the posterior trunk region (**B**, above the elongated yolk) and the CHT **(D)**. **(a’, c’, d’)** are magnifications of regions outlined with white dashed boxes in **(A, C, D)** respectively. Hematopoietic eGFP+ cells were segmented (green contours). White arrows point at *cd34* positive hematopoietic cells. The aorta, the gut, and the sub-aortic clusters are delimited by long blue, magenta, and yellow dashed lines respectively. **(B)** Relative position of eGFP+ cells along the antero-posterior axis of the pronephros (n=279 *cd34*-cells, n=85 *cd34*+ cells). **(E)** Percentage of eGFP+ hematopoietic cells expressing *cd34*, n=6 larvae for the pronephros, n=4 for trunk and CHT regions. **(F)** *cd34* spots counts per hematopoietic cell, n=364, n=705, n=750 cells analyzed for the pronephros, trunk and CHT region respectively. **(B, E, F)** Two sided Wilcoxon tests. NC: notochord, SB: swim-bladder, EY: elongated yolk, G: pronephros glomerulus. Scale bars: 10µm.

Using the *Tg(runx1+23:eGFP)* fish line to perform RNAscope, we observed double positive *cd34*+/eGFP+ cells in the pronephros region in which, as was the case for *gata2b*, they lodge relatively homogenously in comparison to the territory occupied by all eGFP+ cells (**Fig. 7A**, **B**; see also the 3D Imaris reconstitutions **Movie 7**, n=3 larvae at 5dpf). In the AGM and the CHT, double positive *cd34*+/eGFP+ cells were also observed homing more or less inside hematopoietic clusters (**Fig. 7C, D**; see also the 3D Imaris reconstitutions **Movie 8**, n=2 larvae shown). Quantitative analyses revealed that, in both the pronephros and the trunk region, the percentage of *cd34*+ cells among all eGFP+ expressing cells are significantly higher than in the CHT (**Fig. 7E**, with 24.54%, 29.03% and 15.62% for the pronephros, the trunk, and the CHT, respectively). Quantification of RNAscope signals (number of spots/eGFP+ cell, **Fig. 7F**), indicate that the highest signals are detected in double positive *cd34*+/eGFP+ cells localized in the AGM region (trunk), followed by the pronephros and, finally, the CHT. Altogether, these results show that HSPCs are niching in these 3 hematopoietic organs, and that the cells that express the highest amount of *cd34/podocalyxin*, i.e the cells that fit the best the HSC population, are lodging preferentially in the AGM region.

## Discussion

Here, using the zebrafish embryo and larva at early time points of development, we characterize the diversity of developmental HSPCs and progenies as well as of their niches, in the entire organism and at unprecedented spatio-temporal resolution. We have taken advantage of single-cell photoconversion of morphodynamically characterized emerging HSPC precursors, in the ventral floor of the dorsal aorta [described as leading to potential long-term HSCs (Jin et al., 2009; Tian et al., 2017)], to address the potential incidence of emergence heterogeneities on cell fate. We found that the two populations of emerging HSPC precursors – namely EHT pol+ and EHT pol-cells, characterized by their different dynamics and apico-basal polarity features, see (Torcq et al., 2024) –, give birth to multipotent progenies albeit with specific lineage occurrences. This is prominent for the lymphoid lineages, with EHT pol-cells leading more to T-lymphocyte populations and EHT pol+ cells leading more to natural killer populations (NK-like cells). In addition, other remarkable features of EHT pol-cells is the capacity of their daughters to systematically conquer the thymic area, and to differentiate into a population of T-cells among which a significant proportion (22%) expresses *foxp3a*, a functional signature of T-reg cells. Interestingly, the idea of a transient T-reg population produced by precursors emerging from the dorsal aorta, referred to as non-HSC dependent and limited to the developmental period, was raised by Tian et al. (Tian et al., 2017). Of notice, in our EHT pol-cell progenies, we did not detect any potential HSC population that we managed to identify in EHT pol+ cell progenies [this sub-cluster representing approximately 6% of our eHSPC population, was primarily highlighted in our Chromium dataset, with the support of data integration with the Rubin et al adult kidney marrow dataset (Rubin et al., 2022), see **Fig. 5**]. While this could be due to the low sampling of our EHT pol-derived populations, this may also support the idea that these cells are transient, restricted to the developmental period. In this context, we are tempted to speculate that EHT pol-cells would lead to highly potent progenitors that fulfill the apparent multipotency of their progenies, recovered in our MARS-seq dataset. Hence, these should not correspond to the restricted progenitors that have been described in previous studies performed in the zebrafish embryo, such as bipotent T-lymphoid/myeloid progenitors (He et al., 2020), or bipotent lymphoid/erythroid progenitors (Ulloa et al., 2021). These EHT pol-derived, highly potent, progenitors could lead to a sub-population of the eHSPCs that ultimately fails to integrate the niche providing cues necessary to maintain the highest degree of stemness (ex: eHSCs specific of the early developmental period). These may be features more specifically held by the EHT pol+ daughter cells whose eHSPC population contains potential HSCs sharing adult HSC signatures (**Fig. 5H**).

More in line with the cell biological features of EHT pol+ versus EHT pol-cells, i.e apico-basal polarity maintenance of EHT pol+ cells during emergence and not for EHT pol-cells, these may support specific dynamic behavior after their release from the aortic floor (ex: cell migration, intrusion/extrusion throughout vessels etc …) and influencing their ability to conquer more or less specific niches, in particular the ones involved in differentiation and maintenance of long term regenerative potential. In regard to lymphoid lineages, this may be the reason why we observed a significant difference in the thymic areas colonized by EHT pol- and EHT pol+ progenies (with EHT pol+ derived thymocytes being significantly more embedded into the maturing thymus). This difference may also be explained by the nature of the lymphoid cells differentially produced by the two EHT cell types (T-cells and NK-like cells, both having been described to differentiate inside the thymus (Rothenberg et al., 2016)), even if the thymus is not structurally functional at 5dpf.

In this work, throughout the analysis of the entire larval body, we managed to identify relatively highly diversified populations of hematopoietic and immune cells, among which and more particularly several populations of NK-like cells, innate-lymphoid cells (ILC2- and ILC3-like that, for the first time, are detected so early during zebrafish development), and sub-populations of myeloid cells such as early eosinophils. In the latter, we identified a minor fraction expressing the functional marker *eslec* [*si:dkeyp75b4.10* or eosinophil-specific lectin, see (Li et al., 2024)], belonging to the protein family of mammalian *embp* [eosinophil basic major protein, see (Herbert et al., 2024 preprint)], a selectin recently identified as a highly specific eosinophil marker. Interestingly, eosinophils have been shown to undergo a burst of emergence around 5-7dpf in the zebrafish larva (Li et al., 2024), which should explain why we have been able to detect them that early, but the rationale behind this rapid expansion and its functional significance remain unknown. Because, in our datasets, these early eosinophils differentially and significantly express *gata2b* as well as pan-HSPC/MPP markers (such as for example *si:ch211-161c3.6* that is expressed recurrently in most of our developmental HSPC/MPP populations, see **Table S1**), we initially mistook them as a sub-group of eHSPCs. We speculate that these characteristics in expression reflect their very fast engagement in the eosinophil lineage, hypothetically from developmental HSCs and rapidly evolving throughout lineage commitment.

Here we wish to highlight at least one of the intriguing observations made on the eosinophil populations that we describe. In comparison to the markers that we have addressed via *in situ* hybridization using RNAscope (*gata2b, cd34/podocalyxin, cmyb*), some of the cells that strongly express *timp4.2* (an inhibitor of metallopeptidases involved in the degradation of the extracellular matrix, one of the most highly expressed gene of our eosinophil cluster, see **Table S4**) bear the unique feature of accumulating in a sub-region of the pronephros, ahead of the glomerulus. Owing to the described functions of eosinophils, beside innate immune defense, in tissue remodeling by disrupting matrix integrity (Ramirez et al., 2018) and the undergoing maturation of the pronephros region as a niche at 5dpf, it is possible that this population plays specific roles in preparing local niches for hematopoietic cells homing later in the kidney marrow of the juvenile/adult. In addition, our eosinophil cluster expresses many other genes that may relate to niche functions and matrix modulation, such as serine protease inhibitors of the Spink family and several members of the MFAP locus (see **Tables S1, 4**). This may be transposed to the existence of functional heterogeneities in the niches of the adult kidney marrow that will be worth investigating in the future. Finally, the pronephros implantation of cells of the eosinophil lineage may be essential for their differentiation; in the adult zebrafish also, a minor fraction of them is localized in the kidney marrow [while the majority is found in the peritoneal region, see (Balla et al., 2010)].

The markers differentially expressed in cell populations identified in our datasets also allowed, in addition to the pronephros, addressing hematopoietic cell localization in other regions of the larval body, more specifically the CHT and vascular niches in the AGM/trunk region. On this line, we made two fundamental and significant observations.

First, while investigating the localization of *gata2b* expressing cells (eHSPCs and MPPs in majority), we found that, in the upper part of the AGM/trunk region, a minority of hematopoietic cells expressing eGFP driven by the *runx’1+23’* enhancer co-express *gata2b* while a majority of endothelial and endothelial-derived cells (expressing eGFP driven by the *kdrl* promoter) co-express *gata2b*. As visualized in our confocal images, these endothelial cells belong almost exclusively to the SIA, a small artery adjacent to the gut epithelium that is part of the vessels surrounding the gastro-intestinal organs. Of notice, *gata2b* (which is upstream of *runx1*) is also expressed in the dorsal aorta, during the time window of EHT (Butko et al., 2015). Also, *cmyb* and eGFP double positive cells were observed nearby and in close contact with the SIA, indicating that cells of hematopoietic nature, relatively newly born from vessels, are homing in this region. Altogether, these results show that the trunk region, particularly in its upper part, contains at least two niches, including typical sub-aortic *cmyb* positive AGM clusters (Murayama et al., 2006; Zhang and Rodaway, 2007) and, to our knowledge, a not-yet described one located in the direct vicinity of the SIA. We wish to mention here that SIA endothelial cells all along the trunk that are expressing *gata2b* were also visible, although with a lesser frequency than for the upper part of the trunk (not shown). Of notice, the SIA is part of a plexus that includes the sub-intestinal vein and, in the upper part of the trunk, extended inter-connecting vessels (the system establishes around 2 to 4dpf and is delayed in comparison to the early trunk vasculature, see (Isogai et al., 2001)). These different vessels were proposed to initiate from angioblasts homing in the posterior cardinal vein, extruding, migrating, and ultimately contributing to both the arterial and veinous components of the zebrafish peri-intestinal plexus (Goi and Childs, 2016; Hen et al., 2015; Koenig et al., 2016). Owing to its proximity to the gastro-intestinal tractus, the SIA region may be the ideal niche for innate lymphoid cells for example, according to the described functions of innate immune cells of this lineage in the gut (Hernández et al., 2018). Indeed, in our MARS-seq dataset obtained from dissected trunks, we found that ILC2-like cells are enriched in this region. Finally, it also exists the intriguing possibility that the SIA possesses hemogenic capabilities shifted in time in comparison to the dorsal aorta. This idea is strengthened by the fact that at that stage (5dpf), *gata2b* positive cells expressing a high level of eGFP driven by the *kdrl* promoter are virtually never observed contacting the dorsal aorta while it is the case for the SIA, around which we observed round-shaped cells co-expressing eGFP and *gata2b,* with eGFP levels comparable to the ones expressed in contacting SIA endothelial cells (see **Fig. 6g’, g’’**). Finally, in our *Tg(runx1+23:eGFP)* transgenic fish line, we did not find any evidence of eGFP expression in the SIA endothelium at 5dpf, suggesting that *runx1* may not be expressed in this arterial vessel. Hence, if the SIA is indeed hemogenic, it may be one of the sources of hematopoietic stem/progenitor cells supporting the partial recovery of hematopoiesis in *runx1* loss of function zebrafish mutants (Bresciani et al., 2021; Sood et al., 2010).

Second, in entire larvae, we found that cells that express high levels of *cd34/podocalyxin* (*si:dkey-261h17.1* in our sc-RNAseq datasets), hence presumably eHSPCs, localize in the pronephros, the AGM/trunk and the CHT. This is complemented by our sc-RNAseq analyses performed on cells dissociated from dissected body regions in the MARS-seq approach and showing eHSPC clusters present in rostral, trunk and tail regions. Consistently, eHSPCs were also significantly found in the anterior and tail regions, in our Chromium dataset. These localizations, particularly in the AGM/trunk and CHT regions, are consistent with recent results indicating that both regions are establishing functional niches for HSPCs maintenance (Murayama et al., 2023). Interestingly, in whole mount *in situ* hybridizations, many of these cells are not integrated in hematopoietic clusters but appear dispersed and sparse, including in the pronephros region (see 3D Imaris reconstitutions in **Movies 6**, **7**, that allow appreciating the degree of hematopoietic cell compaction). This is reminiscent of what was previously shown in the CHT and in the pronephros, during development of the zebrafish, with relatively sparse HSPCs surrounded by different cellular constituents of niches (Agarwala et al., 2022; Tamplin et al., 2015).

More precisely and based on quantitative analyses (see **Fig. 7E**), we observed that eHSPCs are significantly enriched in the pronephros and in the AGM/trunk region, in comparison to the CHT. In this regard, in our sc-RNA-seq data, cells with the highest differential expression of *cd34/podocalyxin* co-express adult HSC signatures (*nanos1, spi2, meis1b* as well as *gata2b)*, as shown with the integration with the Rubin at al adult kidney marrow dataset [(Rubin et al., 2022), see **Fig. 5G**]. Since the cells in the AGM/trunk region express significantly *cd34/podocalyxin* at the highest average level, they most probably correspond to developmental HSCs (**Fig. 7F**). Altogether with the fact that some of the progenies of single photoconverted cells from the floor of the dorsal aorta remain nearby the site of emergence, this suggests that the AGM/trunk region may be a primary niche for developmental HSC differentiation, from newly emerged pre-HSC. These niches may be, at least in part, the functional equivalent of AGM and/or intra-aortic hematopoietic clusters into which gradually maturing pre-HSCs have been identified in other vertebrate species, including humans (Medvinsky et al., 2011; Rybtsov et al., 2011).

Finally, the specific cell biological features of EHT cell types that we have described in the zebrafish embryo and that fundamentally rely on controlling apico-basal polarity, may or may not be at play in other species. However, if this is the case, they may guide for choosing appropriate tissue culture conditions for *in vitro* reconstitution assays aimed at producing hematopoietic cells with different degree of stemness. Also, the issue of more-or-less selective commitment towards lineages, starting from specific emergence sites, emerging cell types, and continuing with instructions from niches, will be hopefully clarified by highly spatially resolved and unbiased long-term fate mapping strategies associated with single cell and population analyses at different stages and *in toto*, in entire animals. On this line, the developing zebrafish will remain un unvaluable model.

## Materials and Methods

### Contact for reagent and resource sharing

Further information and requests for resources and reagents should be directed to and will be fulfilled by the corresponding Authors, Anne A. Schmidt and Léa Torcq.

### Zebrafish husbandry

Zebrafish (Danio rerio) of the AB background and transgenic fish carrying the following transgenes *Tg(kdrl:nls:kikume)* (Lazic and Scott, 2011); *Tg(cd41:eGFP)* (Lin et al., 2005); *Tg(kdrl:eGFP)* (Jin et al., 2005); *Tg(gata2b:Gal4;UAS:RFP)* (referred to as *Tg(gata2b:RFP*) derived from D. Traver’s *Tg(kdrl:Gal4;UAS:lifeAct-eGFP)* (Butko et al., 2015); *Tg(runx1+23:eGFP)* (this paper) were raised and staged as previously described (Kimmel et al., 1995). Adult fish lines were maintained on a 14 hr light / 10 hr dark cycle. Embryos were collected and raised at 28.5 or 24°C in Volvic source water complemented with 280 µg/L methylene blue (Sigma Aldrich, Cat#: M4159) and N-Phenylthiourea (PTU, Sigma Aldrich, Cat#: P7629) (0.003%, final concentration) to prevent pigmentation. Embryos were all manually dechorionated between 24 and 35hpf. All anesthesia were performed using tricaine methanesulfonate (MS-222, hereafter called tricaine, Sigma-Aldrich Cat#A5040), at a final concentration of 160 μg/ml. Embryos and larvae used for imaging were 2-5dpf, precluding sex determination of the animals. All experiments involving the *Tg(kdrl:nls-kikume)* embryos (including embryo handling, imaging, dissections and dissociations) were performed in the dark without ambient light and with the use of high-pass filters for manipulations under binoculars to avoid accidental photoconversion. All embryos used for single cell RNA-seq experiments were bleached at 10hpf. Briefly, the eggs were bathed in a 0.003% sodium hypochlorite solution in Volvic water for 5 min, washed 3 times 5 min in Volvic water baths, and subsequently raised as described above. The fish maintenance at the Pasteur Institute follows the regulations of the 2010/63 UE European directives and is supervised by the veterinarian office of Myriam Mattei.

### Establishment of the *Tg(runx1+23:eGFP)* fish line

To establish the *Tg(runx1+23:eGFP)* fish line, the pG1/runx1+23-beta-globin-eGFP plasmid was obtained from amplification of the runx1+23-beta-globin and eGFP sequences with overlapping primers and subsequently cloned into the pG1 vector (containing a tol2 inverted terminal repeat), using the Gibson assembly assay (NEB Cat#E2611S). Plasmid was injected at the one cell stage. All steps and materials were performed and obtained according to (Lancino et al., 2018). Importantly, to maximize signal and limit mosaicism, the experiments performed in this work were obtained using the F1 generation.

### In vivo confocal imaging

Embryos were anesthetized using tricaine and positioned in a glass bottom dish (μ-Dish 35mm high; Ibidi, Cat# 81156 or µ-Slide 8 Well high Glass Bottom; Ibidi, Cat# 80807) on their right-side during embedding in 1% low melting agarose (Promega, Cat#V2111) supplemented with final concentrations of 80µg/ml tricaine and 0.0015% PTU in Volvic water. After solidification of the agarose, Volvic water (supplemented with 0.003% PTU and 160µg/ml tricaine) was added to avoid dehydration during acquisitions. Confocal acquisitions were achieved using an Andor (Oxford Instruments) spinning disk confocal system (CSU-W1 Dual camera with 50 μm disk pattern and single laser input (445/488/561/642 nm), LD Quad 405/488/561/640 and triple 445/561/640 dichroic mirrors), equipped with a Leica DMi8 fluorescence inverted microscope, CMOS cameras (Orca Flash 4.0 V2 + [Hamamatsu]) and 63x water immersion objective (HC PL APO 63 x/1.20 W CORR CS2) or 40x water immersion objective (HC PL APO 40x/1.10 Water CORR CS2). The Metamorph software was used for system piloting and image acquisitions.

### Single cell tracing

The transgenic fish line *Tg(kdrl:nls-kikume)* expressing the photoconvertible protein Kikume in endothelial cells was used to photoconvert single EHT-undergoing cells during their emergence from the floor of the dorsal aorta and subsequently trace their progenies in the different developmental and definitive hematopoietic organs. Both EHT pol+ and EHT pol-cells were photoconverted for comparative analysis and discriminated based on morphological characteristics. EHT pol+ cells presented a cup-shape morphology, with an invagination of their luminal membrane. EHT pol-cells displayed a rounded morphology and protruded partially in the aortic lumen. Because there are fewer EHT pol-than EHT pol+ cells, and because of their rounded morphology reminiscent of dividing cells necessitating caution during their selection, out of the 8 independent experiments performed, only 6 included EHT pol-cells and less EHT pol-than EHT pol+ cells were analyzed (n=12 and n=28 respectively). Photoconversions were performed in the trunk region of embryos (along the elongated yolk, between the 7^th^ and 16^th^ inter-segmentary vessels [ISV]) at 2dpf, during the peak time window of EHT. After photoconversion, embryos were raised individually until 5dpf, when progenies of photoconverted cells were traced and imaged in the larvae *in toto*.

#### Single-cell photoconversion and spinning disk confocal acquisition

Single-cell photoconversion was achieved using the aforementioned Andor confocal system with the 63x water immersion objective. UV-illumination at 405 nm (pulses of 2 sec at 25-30% LED power, CoolLED pE-4000 16 wavelength LED fluorescence system) was focused on single EHT-undergoing cells using a Digital Mirror Device (DMD-Mosaic 3, Andor, with the support of the MetaMorph software to manually trace the target region). Z-stack acquisitions at 488 and 561 nm of the region of interest (full camera field of view) were taken before and after photoconversion, to ascertain the specificity of the single-cell photoconversion, using a 408 nm high-pass filter to avoid accidental photoconversion. All acquisitions (at 2 and 5dpf) were taken using identical parameters (objectives, exposure time, laser power) to perform downstream fluorescence intensity measurements and quantitative comparisons.

#### Analysis of EHT progeny migration and multiplication

Cells expressing photoconverted Kikume proteins were manually counted in all hematopoietic organs (the AGM, the CHT, the thymus as well as the pronephros region) using the z-stacks acquired at 5dpf, to compare the distribution of progenies of photoconverted EHT pol+ and pol-cells. At this stage, embryos where accidental photoconversion occurred (during the handling of the embryos, visible by the presence of photoconverted Kikume proteins in vascular structures) were excluded from further analysis. We considered that accidental photoconversion occurred when fluorescence intensity in endothelial cells at 561 nm was above 105% of average background signal. Manual counting was preferred over automatic segmentation for this analysis because of the high variability of fluorescence intensity of progeny cells, of the complexity of the hematopoietic organ structures, especially the vascular plexus of the CHT and the AGM, and of the presence of pigment cells, particularly in the CHT (see asterisks, **Fig. 1F**). Contrary to the pigment cells that vary a lot in size (from 3 µm to 20 µm diameter) and whose auto-fluorescence (at 561 nm) is heterogeneously distributed in the cytoplasm, newly generated hematopoietic cells consistently have a diameter of around 5 µm, a round blast-like morphology and a homogeneous fluorescence pattern. Comparison of the number of progenies per organ as well as of percentage of colonization per organ and per independent experiment was performed using two sided Wilcoxon tests. All quantifications for **Fig. 1C, E, G, I** are available in the **Source Data Table.**

#### Analysis of cell position in thymic region

Spatial analysis of progenies of photoconverted cells was performed using the Imaris software (Oxford Instruments, version 10.1.0). Briefly, progenies of photoconverted cells were segmented using the “Surface” plug-in. Because hematopoietic cells in the thymus still express non-photoconverted Kikume proteins, and because this fluorescent reporter is expressed not only in the nucleus but also the cytoplasm of hematopoietic cells, we extrapolated the exterior surface of the thymus based on the exterior surface of its components (**Fig. 2A** and **Movie 1**). Finally, in each larva we computed the minimal distance between the thymus and each progeny cell. In this case, a negative distance shows the progeny is located inside the thymus while a strictly positive distance shows the progeny is located outside the thymus. For the comparison of cell position relative to the thymus (**Fig. 2C**), cells located more than 15 µm away were considered as completely outside of the thymus and removed from the analysis (thus removing 5 and 6 cells from the total n=52 and n=96 thymic progeny cells of EHT pol-and EHT pol+ cells respectively). This distance of 15 µm is equivalent to 3 times the average thymic cell diameter or 30% of the thymus size (mean diameter of 43 µm). Two sided Wilcoxon tests were used for comparisons. All quantifications for **Fig. 2B, C** are available in the **Source Data Table.** Images of all analyzed thymi are available in **Source data 2.**

#### Analysis of fluorescence dilution post-photoconversion

Undergoing EHT cells as well as their progenies lodging in the thymic area were semi-automatically segmented in Fiji using a custom macro (performing pre-processing, thresholding and size-based segmentation). Fiji was preferred over Imaris for this segmentation step for it allows a better capture of non-convex shapes (such as EHT pol+ cells). Segmentation was followed by manual curation and separation of touching progeny cells using a 3D-watershed (Ollion et al., 2013) whenever necessary. Total fluorescence intensity at 561 nm in the 3D volume of the segmented cell was collected automatically for all cells individually. The total fluorescence intensity in progeny cells was normalized to the total fluorescence intensity of their EHT ancestor for fluorescence dilution analysis. This analysis was performed only on progeny cells lodging in the thymic area, due on the one hand to its circumscribed morphology, enabling whole organ analysis and on the other hand to the biological relevance of this organ for our comparative analysis. Indeed, our results showed clear differences in the capacities of EHT pol+ and pol-cells to generate progenies capable of implanting in the thymus.

We evaluated the validity of our tracing approach, with regards to the stability of the photoconverted Kikume protein as well as the equal repartition of the photoconverted Kikume pool during cell division (**Fig. S1**). For this, single endothelial nuclei (n=4) from the roof of the dorsal aorta in the AGM region were photoconverted at 2dpf (**Fig. S1A, B**) and imaged every day until 5dpf (**Fig. S1A, C**). Cell volumes were automatically segmented as mentioned above and total fluorescence intensity at 561 nm in 3D was collected and normalized, showing a relative stability of the photoconverted Kikume through time (an average of 10% loss at day 3 post-photoconversion, **Fig. S1D**) as well as a comparable fluorescence signal in the daughter cells post-division (2% variability, n=1 endothelial cell out of 4 divided during the 3 days analysis) (**Fig. S1E**). Leveraging this information, we classified the progeny cells located in the thymic area according to their theoretical division history based on their normalized fluorescence intensity (**Fig. 2D, E**). Two sided Wilcoxon tests were used for comparisons (**Fig. 2D**). Cells with a normalized intensity between 0.5 and 0.25 were estimated to have been through 1 division cycle while cells with a normalized intensity between 0.25 and 0.125 were estimated to have been through 2 division cycle, etc… All quantifications for **Fig. 2D, E** and **Fig. S1D, E** are available in the **Source Data Table.**

#### Analysis of implantation of newly generated hematopoietic cells in the AGM plexus post-emergence

To question the origin of newly generated of cells niching in the AGM region, we photoconverted cells (HE cells or EHT undergoing cells or both types) emerging from the floor of the dorsal aorta in either the trunk region (between ISV 7 and 16) or the caudal region (after the end of the elongated yolk). We then assess the ability of their progeny to colonize the AGM or CHT plexus by 5dpf. Photoconversion were performed between 35 and 55hpf, with 1 to 10 cells photoconverted per embryo. Overall, 85 cells in 46 embryos over 11 independent experiments were photoconverted in the trunk region, and 43 cells in 7 embryos over 3 independent experiments in the caudal region. Data in **Fig. 3G** represent percent of photoconverted larvae per experiments with and without AGM or CHT plexus colonization relative to their region of emergence. Two sided Wilcoxon tests were used for comparisons, all quantifications for **Fig. 3G** are available in the **Source Data Table**. For cells emerging in the trunk region, in few embryos, an endothelial cell neighbouring the emerging cell of interest was additionally photoconverted so as to act as a landmark to study the localization of progeny cells in the AGM relative to their region of emergence.

### Single-cell suspension preparation and cell sorting strategies for transcriptomic analysis

#### Cell dissociation

Single cell suspension for FACS sorting was prepared using an optimized protocol adapted from multiple sources (Bresciani et al., 2018; Manoli and Driever, 2012; Samsa et al., 2016). All steps were performed with cooled solutions (4°C), and samples were kept on ice throughout the processing to preserve the viability of cells. Briefly, 5dpf larvae were anesthetized using balneation in embryo medium supplemented with tricaine, at a final concentration of 640 µg/ml. Larvae (between 20 and 30 whole larvae or 60-80 dissected segments per tube) were washed twice in PBS before centrifugation (300 x g for 1 min @4°C) and the supernatant was discarded. Larvae were incubated in 1 ml TrypLE medium (Life Technologies, Cat#: 12605-010) for 10 min on ice, with gentle pipetting every 3 mins, first with a P1000 pipette and subsequently with a P200 pipette. Samples were then centrifugated for 7 min at 300g, before the TrypLE was removed and the cells were resuspended in 500 µl FACSmax medium (Genlantis, Cat#: T200100). Ultimately, samples were passed through a 40 μm cell strainer moistened with FACSmax buffer onto a 35 mm cell culture dish using a syringe plunger. The cell strainer and the dish were washed with 300 µl of FACS max and the flow-through was transferred to a FACS tube and kept on ice until sorting (within the hour).

#### Embryo dissections

For the Chromium (**Fig. 3A**) and MARS-seq (conditions 2, 3 and 4, **Fig. 4A**) strategies, larvae were dissected prior to cell dissociation to collect separately cells from different regions comprising hematopoietic organs of interest. Briefly, larvae were anesthetized using balneation in embryo medium supplemented with tricaine, at a final concentration of 640 µg/ml. Larvae were placed into a petri dish, rinsed in PBS supplemented with tricaine, and a majority of the medium was removed to prevent movement. Larvae were cut using a needle, and were then pooled in tubes and placed on ice until cell dissociation as described above. Two dissection patterns were used. The first one consisted in a separation of the anterior and posterior region of the larvae (2 segments) by a unique transversal cut at the posterior limit of the elongated yolk. For the second pattern, 2 transversal cuts were performed, at the end of the elongated yolk and the posterior limit of the swim bladder, to separate 3 segments: the tail, the trunk and the most rostral region.

#### Cell sorting strategies

Cell sorting was performed on a BD FACS AriaIII cell sorter. For all samples, gating was done on SSC-A vs. FSC-A to collect cells, then on FSC-W vs. FSC-A to keep only singlets and on APC-A to remove dead cells marked by Draq7 (Thermofisher, Cat#: D15106). For the Chromium strategy as well as condition 4 of the MARS-seq, a mix of RFP^+^/eGFP^-^, RFP^+^/ eGFP^med/low^ and RFP^-^/eGFP^med/low^ cells from outcrossed Tg(*cd41:eGFP) X Tg(gata2b:RFP)* larvae were collected, while eGFP^high^ were discarded to prevent specific enrichment of megakaryocytes. For condition 3 of the MARS-seq strategy, eGFP^+^ cells from *Tg(runx1+23:eGFP)* larvae were collected. Finally, for conditions 1 and 2 of the MARS-seq strategy corresponding to the progeny of photoconverted EHT or HE cells using *Tg(kdrl:nls-kikume)* larvae, all cells with photoconverted Kikume (excitable at 561nm) were collected. FACS sorting data were analyzed using either FlowJo (BD), R [flowCore package, (Ellis et al., 2023)] or the open-source flow-cytometry analysis web-tool (https://floreada.io).

### Chromium dataset generation and analysis

#### Transgenic fish line selection

For our dataset construction, that we wanted as exhaustive as possible, we used an outcross of two transgenic reporter lines, Tg(*cd41:eGFP)* and *Tg(gata2b:RFP)*, that both label hematopoietic cell populations. Indeed, our preliminary experiments showed reproducible discrepancies in the labelling of hematopoietic cells by the two reporters (**Fig. S2**). Proportion of single-positive cells vary depending on the region we sample cells from, suggesting that the two transgenic lines label both shared and distinct hematopoietic cell lineages. This discrepancy can be explained by the inherent mosaicism of transgenic lines, the evolution of promoter activation and silencing throughout generations (Goll et al., 2009), and the potential initial incomplete overlap of populations labelled by the two reporters.

#### Library preparation

Based on the 10XGenomics Chromium V3.1 guidelines, a mix of single and double positive cells (**Fig. 3A** and **Fig. S2B**) from approximately 100 dissected 5dpf outcrossed Tg(*cd41:eGFP) X Tg(gata2b:RFP)* double-positives larvae were collected in PBS as described above, and diluted at a concentration of around 400 cells/µL in order to load ∼20 000 cells per condition (anterior and posterior halves of larvae). Chromium Next GEM Single Cell 3ʹ Kit v3.1 (10XGenomics, Cat#: PN-1000269) kit was used, and library preparation was performed according to the manufacturer recommendations (17 PCR cycles were performed for final library amplification, and library quality and concentration were assessed using the BioAnalyzer and Qubit systems). Two independent biological replicates were generated. The sequencing was performed on a Novaseq6000 platform to generate 150-bp paired-end reads, and a theoretical count of 40 000 reads per cell were sequenced (twice the amount recommended by 10XGenomics, to increase chance of observing lowly expressed genes such as transcription factors).

#### Pre-processing

CellRanger (version v6.1.2) was used to demultiplex the reads after sequencing and map them to the zebrafish genome to generate gene expression matrices. For the reference genome, we used the reference genome with improved 3’UTR annotations curated by the Lawson lab [version 4.3.2, (Lawson et al., 2020)], to which we manually added the sequences for RFP and GFP, using the guidelines provided by 10XGenomics for the use of personal modified reference genomes. The use of a genome with enhanced annotations compared to the Ensembl zebrafish genome danRer11 (GRCz11) allowed us to improve median mapped gene per cell count (going from 2139±250 to 2666±297). Datasets were analyzed using R (version 4.2.3) and a custom pre-processing pipeline, using the Seurat package [version 4.1.1, (Stuart et al., 2019)] as well as multiple independent packages, cited below. Pre-processing (low-quality cell removal as well as doublet analysis, clustering and manual annotation) was performed independently on each dataset before merging for final analysis. First, filtering of low-quality cells was performed and thresholds were manually selected based on distribution of gene relative to UMI counts and mitochondrial gene percentage. A minimal threshold of 800 genes per cell and maximal percentage of 5% of mitochondrial genes were ultimately used for all datasets. In addition, cells with abnormal gene to UMI ratios, likely corresponding to either low-quality cells or potential multiplets, were removed (Morizet et al., 2024). An average of 29.3±5.1% of cells were removed at this stage. Counts were normalized (LogNormalize) and scaled before dimensionality reduction (PCA), Principal Components (PCs) selection and clustering. For PCs selection, we kept PCs explaining more than 2% of the variance across the first 100 components and that were significant according to the Jackstraw procedure, ultimately selecting between 45 to 56 PCs for our 4 datasets. Clustering was performed using the Louvain algorithm, with a resolution of 1.2. This preliminary clustering was used to perform a doublet labelling step, using both the DoubletFinder (McGinnis et al., 2019) and the scran DoubletCells (Lun et al., 2016) algorithm. Although these two methods flagged potential doublet-cells, they did not identify any potential artefactual doublet clusters that would impact our clustering, and thus all cells were kept for further analysis.

#### Analysis

At this stage and before the integration of our 4 datasets, refined clustering and manual annotations was performed independently for all datasets. Clustering was performed at multiple resolutions (0.3-2), and the stability and conservativeness of the clustering across multiple resolutive scale was assessed using the Clustree package (Zappia and Oshlack, 2018). Final clustering resolution was chosen based on pairwise comparison of differentially expressed genes (DEG, non-parametric Wilcoxon rank sum test) across neighbouring clusters. Cells identified as contaminants (muscular cells, skin epithelial cells or neural cells) were removed from the dataset at this stage. Finally, all identified clusters were manually annotated so as to evaluate the cell diversity in each dataset and guide our integrating algorithm selection. Indeed, because of the discrepancy observed in term of cell types and prevalence between our regionally distinct datasets, and to preserve after integration the biological disparities observed in our separate datasets, we used for this step the Rliger algorithm (Welch et al., 2019), that uses integrative non-negative matrix factorization and allows to both identify common denominators but also is also conservative of dataset specific factors. The integration steps, including normalization, scaling, integration and quantile normalization was performed as recommended by the developers, using the Seurat wrappers. Finally, clustering and manual annotation was performed on integrated data as described above for separate datasets. Cell type identity was manually assigned based on the Differentially Expressed Genes (DEG, tested with the Seurat FindAllMarkers function, using non-parametric Wilcoxon rank sum test) for each cluster (**Table S1**) and comparison to known lineage-specific markers (**Table S2**). Multiple clusters sharing expression of lineage-specific markers were grouped together into a same cell type identity to facilitate visualization and further analysis (for example clusters 18-21-34-37-40 were all identified as NK-like cells and clusters 16-20-31 were identified as T-Lymphocyte Progenitors, **Table S1**). In such cases, the sub-clustering information was kept for more in-depth analysis of cellular diversity (**Fig. S3**). Dataset visualization was performed using UMAPs with default settings for all plots. Gene expression was visualized by dot-plots or violin-plots. Gene selected for visualization by dot-or violin-plots were identified within the DEG lists or manually curated specific markers of interest. Cell cycle analysis was performed using the Seurat CellCycleScoring function.

### MARS-seq dataset

The MARS-seq protocol (Jaitin et al., 2014; Keren-Shaul et al., 2019) is based on single-cell RNA-sequencing of FACS sorted indexed single cells. This system allows for the combined acquisition of transcriptomic and indexing data of rare cell populations, that can be collected and conserved before parallelized transcriptomic library construction. This method is particularly adapted to be combined to our low-throughput single-cell photoconversion protocol, and allows for the accumulation of cells over multiple photoconversion experiments as well as the integration of the multiple conditions that we detail below.

#### Single-cell photoconversion

For analysis of transcriptomic identity of progenies of EHT pol- and EHT pol+ cells, single-cell RNA-sequencing was performed using the MARS-seq protocol (condition 1, **Fig. 4A**). For this assay, using *Tg(kdrl:nls-kikume)* embryos, single EHT undergoing cells located in the trunk region (between ISV 7 and 16) were photoconverted at the peak time window of emergence (between 48hpf to 58hpf). Embryos were raised separately in the dark until cell dissociation for indexed-FACS in plates at 5dpf. 6 independent experiments were performed, tallying a total of 264 photoconverted EHT pol+ cells and 115 EHT pol-cells, with an average of 44.0±7.0 and 25.2±7.5 cells per experiments respectively.

#### Whole hemogenic endothelium photoconversion

To facilitate the analysis of progenies of single-photoconverted cells (a high specificity but low-throughput strategy), we complemented this first approach by adding in our MARS-seq dataset progenies of whole hemogenic endothelial cells, thus increasing the number of cells for analysis. We performed photoconversions of multiple hemogenic cells (3 to 10 cells per embryo) in the floor of the dorsal aorta in the trunk region (either from ISV 2 to 5 or 7 to 16) of 48-58hpf embryos (condition 2, **Fig. 4A**). Embryos were then raised in the dark until dissection and cell dissociation for indexed-FACS in plates at 5dpf. 9 independent experiments were performed, with an average of 184.1±46.0 photoconverted cells per session. Taking advantage of the higher number of cells generated through this approach, we added an additional layer of spatial information to our dataset by performing larvae dissection before cell dissociation. For the dissection, 3 different modalities were used (see above): no dissection (2 replicates), 2 segment dissection (5 replicates) and 3 segment dissection (2 replicates).

#### Hematopoietic reporter lines

To facilitate the integrative analysis of our MARS-seq and Chromium datasets, we included in our MARS-seq dataset cells collected from transgenic reporter lines, namely from the *Tg(runx1+23:eGFP)* and the outcross of the *Tg(cd41:eGFP)* and *Tg(gata2b:RFP)* lines (condition 3 and 4 respectively, **Fig. 4A**). Cells were collected from 5dpf larvae after dissection (3 segments *Tg(runx1+23:eGFP)* and 2 segments for *Tg(cd41:eGFP)* X *Tg(gata2b:RFP))*. 3 and 2 independent experiments were performed, respectively.

#### Library preparation

Capture plates (384 wells plates) containing a lysis mix (nuclease-free water, 10% Triton X-100, RNasin plus 40U/mL (Promega, Cat#: N2611) and MARS-seq barcodes (Keren-Shaul et al., 2019) including a T7 RNA polymerase promoter, a partial Illumina paired-end primer sequence, a cell barcode followed by a unique molecular identifier (UMI), and a polydT stretch) were prepared in advance using the Bravo automated liquid handling platform (Agilent) and stored at -80°C. Plates were thawed at 4°C and centrifugated before being used for cell collection. Cells were index-sorted into the plates using a FACS AriaIII cell sorter, and the plates centrifuged at 4°C and immediately placed on dry ice before storing at -80°C until processing.

Overall, 28 plates were accumulated and processed during two independent session (14 plates per session), tallying 10404 sorted cells (the total amount of sorted cells per replicate and condition can be found in **Table S3**). Plates were processed according to the MARS-seq2.0 protocol. Briefly, a first step of reverse transcription (RT) (SuperScript III, Thermo Fisher, Cat#: 18080085) was conducted to barcode the transcripts with unique cell and UMI barcodes. After RT, primers were removed with an exonuclease (ExoI NEB, Cat#: M0293L), and the barcoded cDNAs for each single cell in a half-plate were pooled together for the next steps. cDNA was converted to dsDNA (Second Strand synthesis kit, NEB, Cat#: E611L) and linearly amplified by in vitro transcription (HiScribe T7 High Yield RNA Synthesis Kit, NEB, Cat#: E2040S). The dsDNA was degraded (TURBO DNase Fisher, Cat#: 10646175) and the amplified RNA fragmented for sequencing (Fisher, Cat#: 10426914). A pool barcode with Illumina adapters for each half-plate was added by RNA-DNA ligation before a second RT step (AffinityScript Multiple Temperature reverse transcriptase, Agilent, Cat#: 600109). Final amplification by PCR was performed (KAPA HiFi HotStart ReadyMix, Roche Diagnostics, Cat#: 7958935001). The resulting libraries were controlled by qPCR amplification of housekeeping genes (*ef1a* and *ß-actin*), DNA quality control using a TapeStation, Agilent) and DNA concentration using the Qubit dsDNA high-sensitivity assay. Libraries were then sequenced using the NextSeq 500/550 High Output Kit v2 (75 cycles) (Illumina), aiming for a theoretical average of 75000 50bp reads per cell.

#### Pre-processing and analysis

Raw reads were converted to fastq files using bcl2fastq package (2.20.0). Reads were demultiplexed and mapped to the Lawson Lab curated genome [(Lawson et al., 2020), v4.3.2] using the MARS-seq pipeline scripts (Keren-Shaul et al., 2019). Analysis was done using the Seurat package. Manually curated metadata (containing information on transgenic line and regional origin) was added to the Seurat objects. Quality-check, filtering and preliminary analysis were performed independently on each of the 56 libraries, that correspond to the 28×2 half-plates, with identical procedure to the Chromium dataset. All libraries were merged together without a specific integration step for further processing. Indeed, all 56 libraries were inherently composed of cells from different origins (mixes of different transgenic lines and spatial origin), and good merging was obtained without requiring any specific batch-removal steps. Normalization, scaling, dimensionality reduction, clustering and manual annotations were performed as described above for the Chromium dataset (**Table S2, S4**).

### Integration of MARS-seq and chromium datasets

To compare the populations identified in our two datasets, we integrated them together using the Seurat FindIntegrationAnchors and IntegrateData functions with default parameters. Following integration, dimensionality reduction, clustering and differential gene expression analysis was performed as described for the Chromium datasets. Verification of the stability and robustness of our clustering before and after integration was assessed using alluvial plot (**Fig. S5B**), that showed good conservation of cell type identity annotation. Overall, the global architecture of our two datasets and the merge dataset is conserved, as shown in the UMAP visualizations (**Fig. 3B, 4B, S5A**). The identified clusters and sub-clusters were also highly similar based on gene expression analysis, validating the biological relevance of our populations identified across different technical modalities.

### Distribution of cell types relative to larval region and transgenic origin

To compare the distribution of cell types across the different larval region investigated (**Fig. 3E**, **S6A**, **S7A**) or between two conditions (**Fig. 4E**), the percentage of each cell type per replicate was computed and compared when number of replicates >2 (two sided Wilcoxon tests with Holms-adjustments in case of multiple comparisons). Similar method was used to compare cell-cycle state across larval region or between different cell types (**Figs 3F**, **S4A, S6B**, **S7B**). All tables recapitulating distribution of cell type relative to condition (transgenic lines vs photoconverted hemogenic cells progenies and EHT pol+ vs EHT pol-) or region (anterior, rostral, trunk, tail) for MARS-seq and Chromium datasets are available in the **Source Data Table**. Similarly, tables recapitulating cell cycle states percentage relative to region, cell type and/or condition are available in the **Source Data Table**.

### Comparison of developmental and juvenile / adult lymphoid populations

To gain insight on our T-Lymphocyte Progenitors population as well as the closely related innate lymphoid cells (NK-like and ILC-like) and dendritic cells, we downloaded a single-cell RNA-seq dataset of juvenile (4 weeks) and adult (3-5 months) thymic populations (Rubin et al., 2022) and integrated our aforementioned populations to their T-cell, NK, ILC and dendritic cell populations. For the integration we used the Seurat FindIntegrationAnchors and IntegrateData functions, modifying the default parameters to limits the number of selected features to 1500 and regress out variations due to number of genes detected and percentage of mitochondrial genes. After integration, we performed scaling, dimensionality reduction (PCA), Principal Components (30 PCs) selection and clustering (Louvain algorithm). We performed the clustering at various resolutions, from 0.5 to 1.5, so as to be able to re-identify the sub-clusters characterized by Rubin and colleagues. DEG were computed for all clusters, and DEG between developmental and juvenile / adult cells per clusters were also calculated to identify conserved and specific genes expressed in our different populations throughout the different stages analyzed. Because virtually no T-lymphocyte specific genes are expressed in our developmental dataset (*tcr α / β / γ / δ* or *cd4-cd8* surface markers) and were thus not taken into account during the integration steps, non-T-cell populations of our dataset (ILC2, ILC3 and NK-like cells) co-clustered with adult T-cell populations. They could not be discriminated on the basis of the key aforementioned T-cell genes, but on the contrary, they clustered together based on the expression of other immune cell specific genes shared between these closely related populations.

### Comparison of HSPCs population across multiple datasets

To complement our analysis of developmental HSPCs and MPPs populations and compare them to adult definitive HSCs, we downloaded a publicly available single-cell RNA-seq datasets of zebrafish hematopoietic populations (Rubin et al., 2022) for comparison. This dataset was generated using whole kidney marrow cells sorted on side and forward scatter to enrich in granulocytic as well as lymphoid and progenitor fractions. We integrated together our HSPCs and MPPs populations with their multiple HSPCs clusters, as described above for the lymphoid populations (selecting 3000 variable features). After integration, we performed scaling, dimensionality reduction (PCA), Principal Components (23 PCs explaining 2% of the variance) selection and clustering (Louvain algorithm, resolution 0.2). The clustering generated 7 clusters, to which we manually assigned identities based on gene expression (**Table S6**). Qualitative analysis of the repartition of cells in newly defined clusters depending on their origin was visualized using an alluvial plot. For the two most undifferentiated clusters (HSCs and MPPs), we performed DEG analysis to identify genes potentially labelling long-term HSCs (**Table S6, Fig. S8B**). Within the HSCs populations, we compared gene expression between cells coming from our 5dpf dataset and cells coming from adult kidney marrow (**Table S6, Fig. S8C**), and did not find any genes that would be specific of the two stages we compared. DEG analysis was performed using FindMarker functions in Seurat (non-parametric Wilcoxon rank sum test). The expression of genes we identified as being specific of the HSCs cluster was then investigated in the MARS-seq dataset, within the progenies of EHT pol- and EHT pol+. Similarly, the expression of these genes was investigated in other published embryonic hematopoietic populations datasets (Ulloa et al., 2021; Xia et al., 2021).

### In situ gene expression analysis

#### Whole mount single molecule fluorescent in situ hybridization using RNAscope

For RNAscope experiments, we used the *Tg(kdrl:eGFP)* and *Tg(Runx’1+23’:eGFP)* fish lines that allowed localizing RNAscope signals in/around hematopoietic and endothelial cells, and vessels. The GFP fluorescence was partly maintained after fixation, which prevented from performing an immunofluorescence after the RNAscope procedure.

RNAscope was performed on 5dpf larvae as described in (Torcq et al., 2024) with the following modifications. 5dpf larvae were fixed in 4% formaldehyde (Electron Microscopy Sciences, Cat# 15712) diluted in PBS/0.1% tween20 (PBST) for 2hrs at room temperature, washed in PBT and kept in MeOH at -20°C until use. After re-hydration, larvae were incubated with H2O2 for 10 min at RT, washed for 15 min in PBST, incubated for 45 min at 37°C with proteinase K in PBST (1/2000 from a glycerol stock at 20mg/ml, Ambion Cat# 10259184) and washed in PBST for 2×10 min at RT. In subsequent steps of the RNAscope procedure, the Multiplex Fluorescent Reagent Kit v2 was used (ACD Biotechne Cat# 323100), including H2O2; probe diluent (PD); wash buffer (WB); AMP1, AMP2, AMP3 buffers; HRP-C1, HRP-C2, HRP-C3 reagents; TSA buffer; HRP blocker. All incubations were performed in Eppendorf tubes, with a maximum of 20 larvae/tube, in a dry heating block. Before addition of probes, larvae were incubated for at least 2 hrs at 40°C in PD. Larvae were then incubated overnight at 40°C with the *gata2b* (ACD Biotechne Cat# 551191-C2), *cd34* (ACD Biotechne Cat# 1223761-C1, 1223761-C2), *timp4.2* (ACD Biotechne Cat# 1255031-C3), *cmyb* (ACD Biotechne Cat# 558291 (-C1), 558291-C2, 558291-C3) RNAscope probes.

Next morning, larvae were washed for 2×10 min in WB and incubated sequentially with AMP buffers during 30 min for AMP1 and AMP2 and 15 min for AMP3, at 40°C, with 10 min washes at RT between each AMP buffer incubation. Larvae were then incubated for 15 min at 40°C with RNAscope Multiplex FL v2 HRP-C1, HRP-C2, HRP-C3, washed for 10 min at RT with TSA buffer, incubated for 30 min at 40°C with OPAL-570 (Akoya Biosciences Cat# PNFP1488001KT) in TSA buffer (dilution 1/500), washed 2×10 min at RT in WB, incubated for 15 min at 40°C in HRP blocker and finally washed 2×10 min in WB and 2×10 min in PBST at RT. Larvae were then imaged by confocal microscopy.

#### Confocal imaging for Whole mount single molecule fluorescent

RNAscope experiments were imaged similarly to live sample (see *in vivo* confocal imaging paragraph above). Larvae were embedded in 1% low-melting agarose (Promega, Cat# V2111) in PBS 1 X in a glass bottom 60μ-Dish (35 mm high; Ibidi, Cat# 81156). Images were acquired with two confocal microscope systems: either the Andor system described in the “*In vivo* confocal imaging” paragraph above, or a Nikon Ti2e spinning disk microscope, equipped with a sCMOS camera (Photometrics, Prime 95B, pixel size 11 µm, 1,200×1,200 pixels, QE 95%) and a 40x water objective (Numerical Aperture 1.15, Working Distance 0.6 mm, xy pixel size of 0.27 µm). Briefly, the entirety of the larvae was manually scanned to localize cells positive for our genes of interest (GOI). Z-stacks were acquired in 3 major larval regions where hematopoietic cells niche: the pronephros, the trunk region (starting from the end of the swim bladder and ending at the urogenital opening) and the CHT region (starting from the urogenital opening to the end of the posterior cardinal vein). Between 2 and 3 z-stacks per larvae were acquired for the trunk and CHT region, to capture the entirety of the hematopoietic niches.

#### Quantitative image analysis of in situ gene expression data

mRNA expression quantitative analysis was performed using the Imaris software (Oxford Instruments, version 10.1.0). All images were processed similarly to analyze the repartition of GOI positive hematopoietic cells labelled by the Tg(*runx1+23:eGFP)* transgene throughout the different hematopoietic organs. Both cell volume (GFP signal) and RNAscope spots were simultaneously and automatically segmented using the “Cells” tool from Imaris, that segments individual cells (generates an outer surface) as well as RNAscope signal within cells. Automatic segmentation was manually curated, mostly to remove artefactual cells detected within the notochord due to its autofluorescence and its high ability to take up the OPAL dye. For the *cd34* experiments, because of the strong *cd34* RNAscope signal generating fluorescence leakage in the GFP channel, manual curation was also required to remove artefactual cells (cells with virtually 100% overlap between the two GFP and RNAscope channels). Finally, all quantitative information, including hematopoietic cell count, GOI positive cell count, positional information and number of spots per cell were retrieved from the ‘Statistics’ tab in Imaris. Cell count and spot count per cell were used to compare proportion of positive cell per region (**Figs 6F, 7E**, **S9E**) as well as number of spots per cell per region (**Fig. 7F**). The data plotting and statistical analysis was performed using R software.

For *gata2b*, we analyzed n=3 whole 5dpf larvae with additional n=12 pronephros (total n=15) from *Tg(runx1+23:eGFP*) background (**Fig. 6B, F**), n=4 anterior trunk regions from *Tg(runx1+23:eGFP*) background (**Fig. 6H**) and n=6 anterior trunk regions from *Tg(kdrl:eGFP*) background (**Fig. 6H, J**).

For *cmyb*, we analyzed n=4 anterior trunk regions of 5dpf larvae from *Tg(kdrl:eGFP*) background (**Fig. 6K**)

For *timp4.2*, we analyzed n=3 whole 5dpf larvae with additional n=6 pronephros (total n=9) from *Tg(runx1+23:eGFP*) background (**Fig. S9B, E**).

For *cd34*, we analyzed n=4 whole 5dpf larvae with additional n=2 pronephros (total n=6) from *Tg(runx1+23:eGFP*) background (**Fig. 7B, E**), within which 364, 705 and 750 cells hematopoietic cells localized in the pronephros, trunk and tail region respectively were compared, **Fig. 7F**).

For the analysis of GOI positive cells position within the pronephros region (for *gata2b*, *cd34* and *timp4.2* respectively, see **Figs 6B, 7B**, **S9B**), spatial positions of all hematopoietic cells (GFP+) were recovered and normalized per larva to allow inter-individual comparison. Briefly, each cell position along the antero-posterior axis of the larva (x-axis) was normalized according to the following formula (z-score normalization):

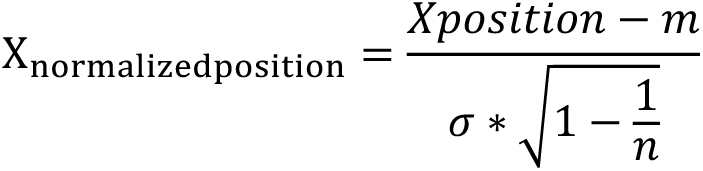

with *Xposition* the position along the x-axis, *m* the mean of the sample, σ the standard deviation of the sample and *n* the number of observations, the sample being all the GFP+ cells (GOI+/-) in a single larva.

To compare the expression of *gata2b* in hematopoietic and vascular cells [labelled by the transgenic lines *Tg(runx1+23:eGFP)* and *Tg(kdrl:eGFP)* respectively] in the upper trunk region around the SIA, RNAscope spots were segmented independently of GFP+ cells (using the Imaris Spots function). This allowed to evaluate the percentage of *gata2b* spots per larva that localized in and out of hematopoietic or vascular cells (see **Fig. 6H**).

To compare the expression of *gata2b* and *cmyb* in vascular and hematopoietic cells (labelled by the *Tg(kdrl:eGFP)* line, see **Fig. 6J, K**), GFP+ cells were segmented (using the Cells tool from Imaris) and manually classified into 3 categories, based on their proximity to the 3 main vascular structures visible in the upper trunk region: the dorsal aorta, the vein plexus below the aorta and the SIA. Cells that could not be associated with any of those structure (located farther than 10µm from any structure) were not classified and removed from further analysis.

All quantifications for **Figs 6B**, **F, H, J, K, 7B, E, F, S9B, E** are available in the **Source Data Table.** Two sided Wilcoxon tests were used for comparisons, except for data relative to **Fig. 7F** were Welch’s t-test with Holm’s adjustment were used.

## Supplementary Figure Legends

**Supplementary Figure 1:**
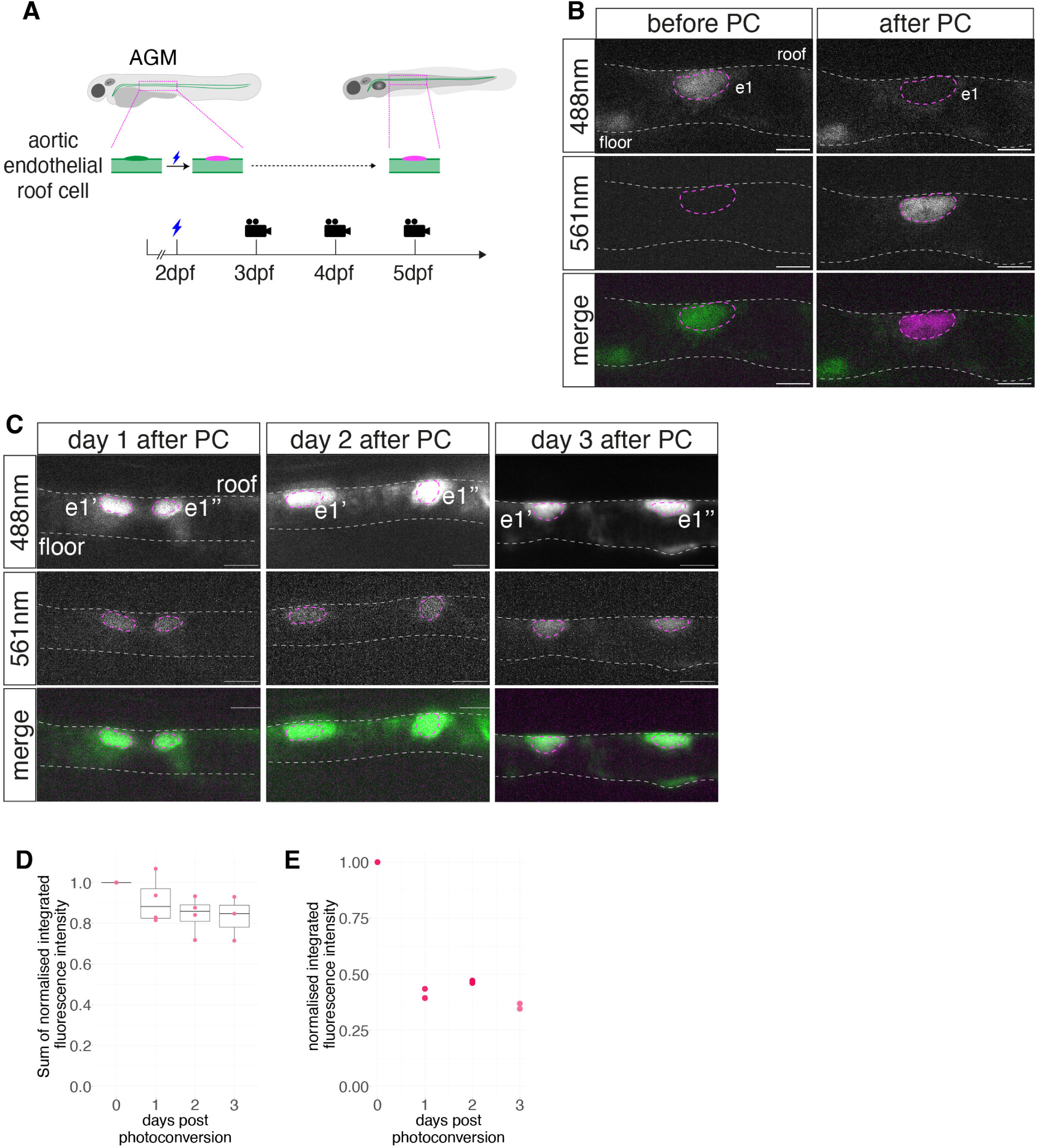
The photoconverted Kikume protein is stable and partitioned equally into daughter cells after division. **(A)** Strategy for single endothelial cell photoconversion. Single endothelial cells of the *Tg(kdrl:nls-kikume)* fish line at 2dpf, located on the roof of the dorsal aorta in the AGM region, were photoconverted so as to study the stability and dilution of the photoconverted Kikume protein. This strategy was used as a reference for our single-EHT photoconversion strategy, as endothelial cells do not migrate or divide extensively at this stage. A unique endothelial cell was photoconverted at 2dpf in each embryo (n=4), that were subsequently raised separately and imaged every day, for 3 days. **(B)** Representative spinning disk confocal images (maximum z-projection) of a single aortic endothelial cell (e1, outlined in magenta) before and after photoconversion (PC) (left and right panels respectively). The cell before photoconversion can be visualized at 488nm only. After photoconversion, the targeted cell is visualized at 561nm and most of the fluorescence at 488nm has been extinguished. **(C)** Spinning disk confocal images (maximum z-projection) of the two daughters of a photoconverted endothelial cell, (e1’ and e1’’, outlined in magenta), resulting from the division of cell e1 in panel **(B)**. The fluorescence at 561nm is quite stable over time and comparable in both daughter cells. The fluorescence at 488nm increases over time due to the maintained activity of the *kdrl* promoter in endothelial cells, thus resulting in neo-synthesis of Kikume protein. **(B, C)** White dashed lines: aortic roof and floor. Scale bars: 10µm. **(D)** Evolution of normalized fluorescence intensity at 561nm over time. n=4 single endothelial cell photoconversion experiments. Total integrated intensity of fluorescence at days 1 to 3 was normalized to integrated intensity of fluorescence immediately post-photoconversion. **(E)** Evolution of normalized fluorescence intensity at 561nm in 2 daughter cells over time (n=1). Total integrated intensity of fluorescence of daughter cells at days 1 to 3 was normalized to integrated intensity of fluorescence of the mother cell immediately post-photoconversion.

**Supplementary Figure 2:**
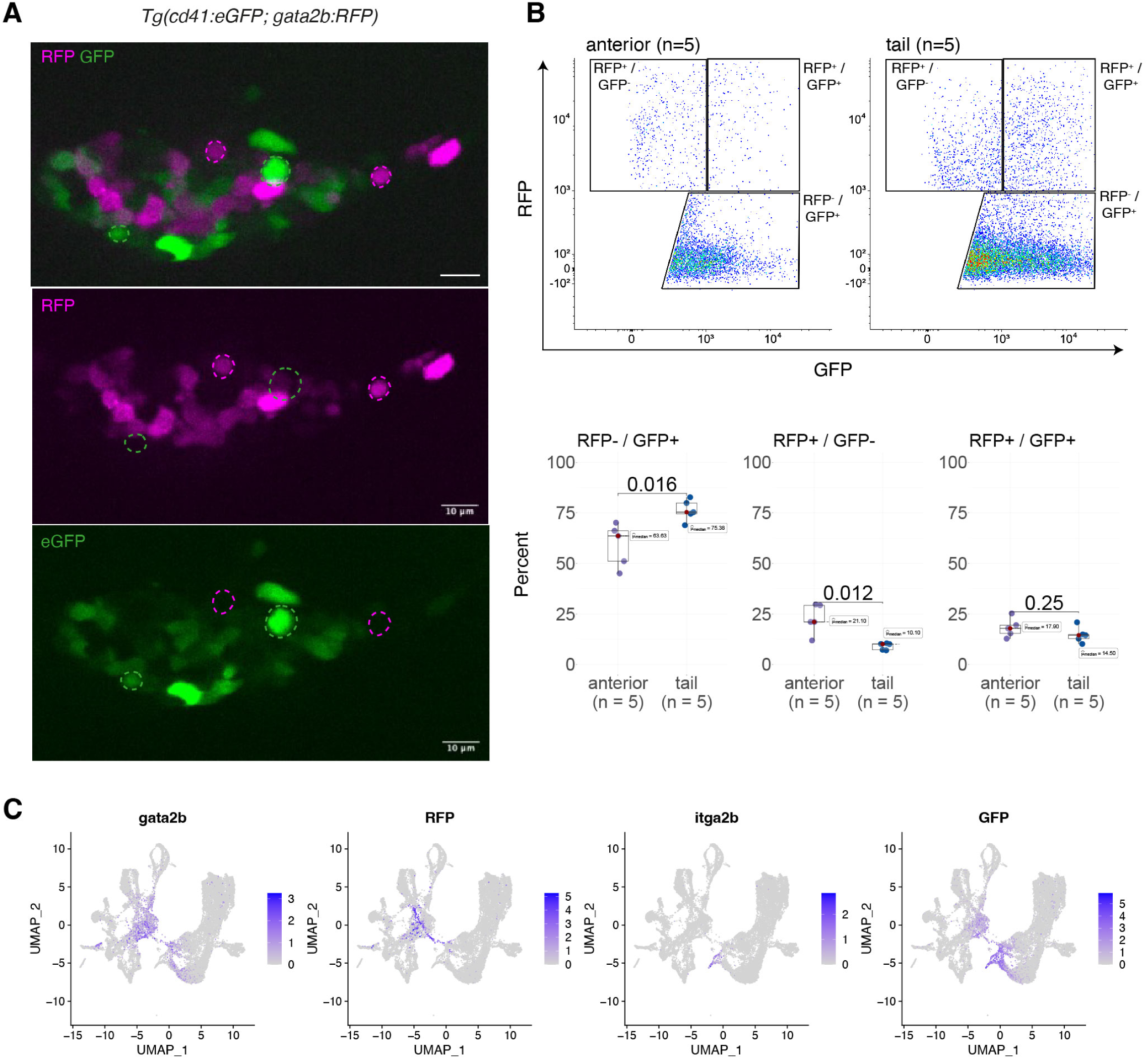
Reference transgenic lines labelling hematopoietic cells are incompletely overlapping. **(A)** Representative spinning disk confocal images (maximum z projection) of a thymus of a double transgenic *Tg(cd41:eGFP; gata2b:RFP)* 5dpf larva. Cells outlined in green or magenta are single-positive cells either for eGFP or for RFP respectively. Scale bar: 10µm. **(B)** Top panels: representative FACS gating strategy for *Tg(cd41:eGFP; gata2b:RFP)* cell sorting at 5dpf from anterior and tail regions. Gates delimit the single and double positive populations, that were ultimately sorted together for transcriptomic analysis. Bottom panels: Comparison of proportion of double and single positive populations in anterior and posterior region (n=5 replicates), showing that RFP single positive cells are over-represented in the anterior region compared to the tail, while GFP single positive cells are over-represented in the tail compared to the anterior region (two-sided Wilcoxon tests). **(C)** UMAPs of *gata2b* and *itga2b* (*cd41*) mRNA expression as well as of our two fluorescent reporters *rfp* and *gfp,* in our Chromium dataset. This data shows that although cells were sorted based on the presence of a fluorescent protein expressed under the control of hematopoietic promoters, the expression of fluorescent reporter mRNAs is restricted to a fraction of the cells, owing to the different half-life, synthesis rate and detection limit of mRNAs and proteins. Also, the expression of the fluorescent reporter mRNA does not match exactly the expression of their hematopoietic reporter counterpart (*eGFP/itga2b*, *RFP/gata2b*), highlighting a potential mismatch between the activation of the promoter of the transgene and the endogenous promoter.

**Supplementary Figure 3:**
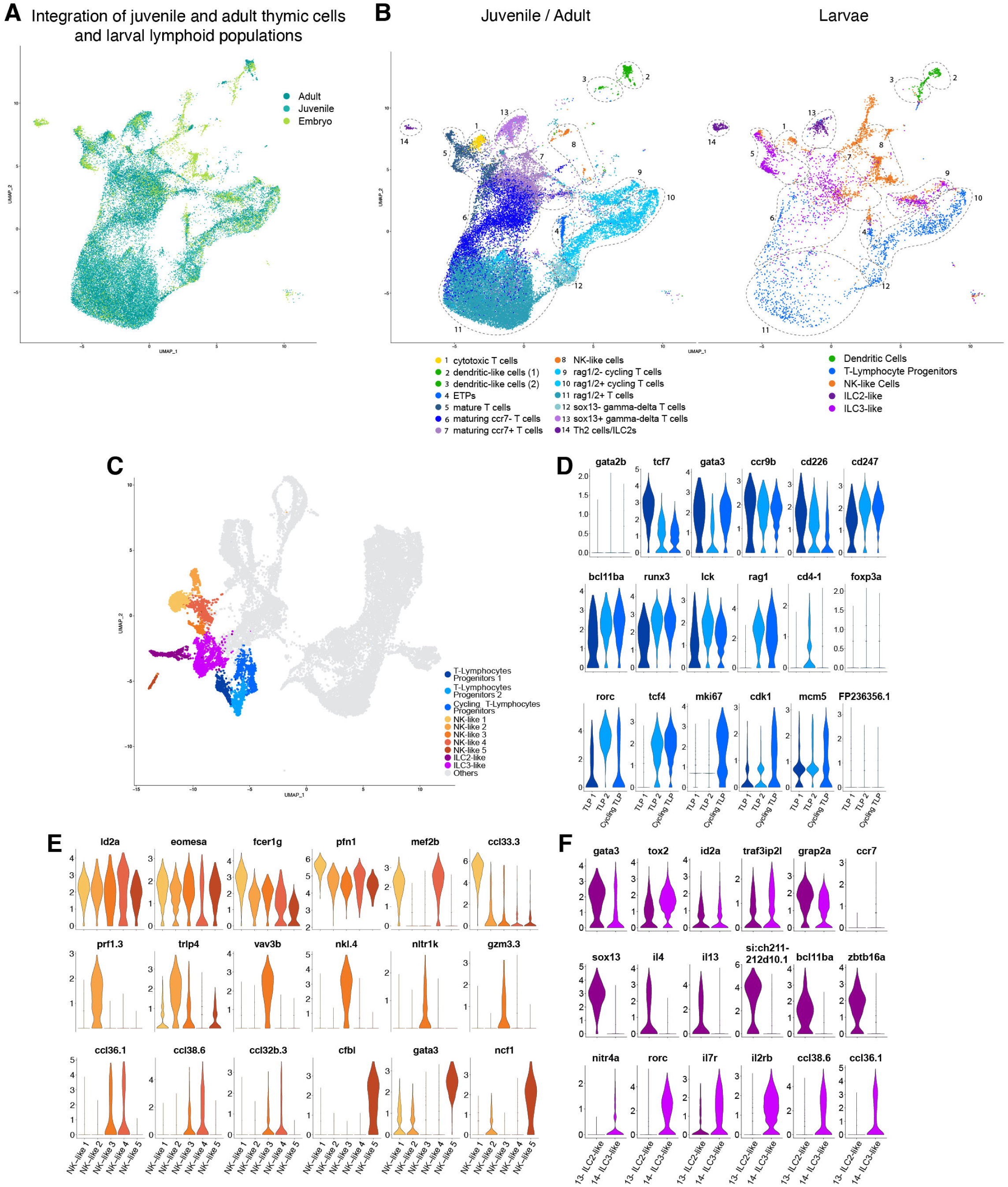
Integrated analysis of larval, juvenile and adult lymphoid populations. **(A)** UMAP of integrated lymphoid populations from different stages: larval (clusters 10 to 14 of our Chromium dataset, generated with double and single positive cells from 5dpf whole *Tg(cd41:eGFP; gata2b:RFP)* larvae), juvenile and adult [whole thymic cells from 4wpf and 3-5mpf zebrafish, (Rubin et al., 2022)]. The UMAP is colored by stage. **(B)** UMAPs of integrated lymphoid populations, split by stage (left: juvenile/adult, right: larval). Clusters are colored by their identity (our manual annotation for the developmental dataset, original published annotations for the juvenile/adult dataset). Larval ILC2-like population clusters both with the adult Th2 Cells/ILC2 (#14) and the sox13+ gamma-delta T cells (#13), with which, for the latter, they share the expression of *sox13*. Larval ILC3-like population clusters in majority with mature T Cells (#5), maturing *ccr7*+ T Cells (#7) and *rag1/2*-cycling T-cells (#9), owing to the shared expression of *ccl38.6/tnfs14/syk, ccr7/il7r/id3* and cell cycle markers, respectively. Larval NK-like Cell populations cluster with NK-like Cells (#8), cytotoxic T Cells (#1), ETPs (#4) and *rag1/2*-cycling T-cells (#9). Clustering with cytotoxic T-cells is due to genes such as granzymes or *eomesa*, while clustering with rag1/2-cycling T-cells is based on shared expression of cycling genes and absence of *rag1/2* expression. Clustering with ETPs can be explained by the shared expression of *gata3*. Larval T-Lymphocyte progenitors (TLP) population clusters with ETPs (#4), *rag1/2*+ cycling T-cells (#10), and *rag1/2*+ T-cells (#11), owing to the shared expression of *gata3/hells/notch1b*, cell cycle markers and *rag1/rorc/lck/ccr9b/bcl11ba/runx3* respectively. **(C)** UMAP of lymphoid lineage subpopulations found in our Chromium dataset. **(D, E, F)** Violin plots of differentially expressed genes (non-parametric Wilcoxon rank sum test) specific of TLP, NK-like and ILC-like sub-populations, respectively.

**Supplementary Figure 4:**
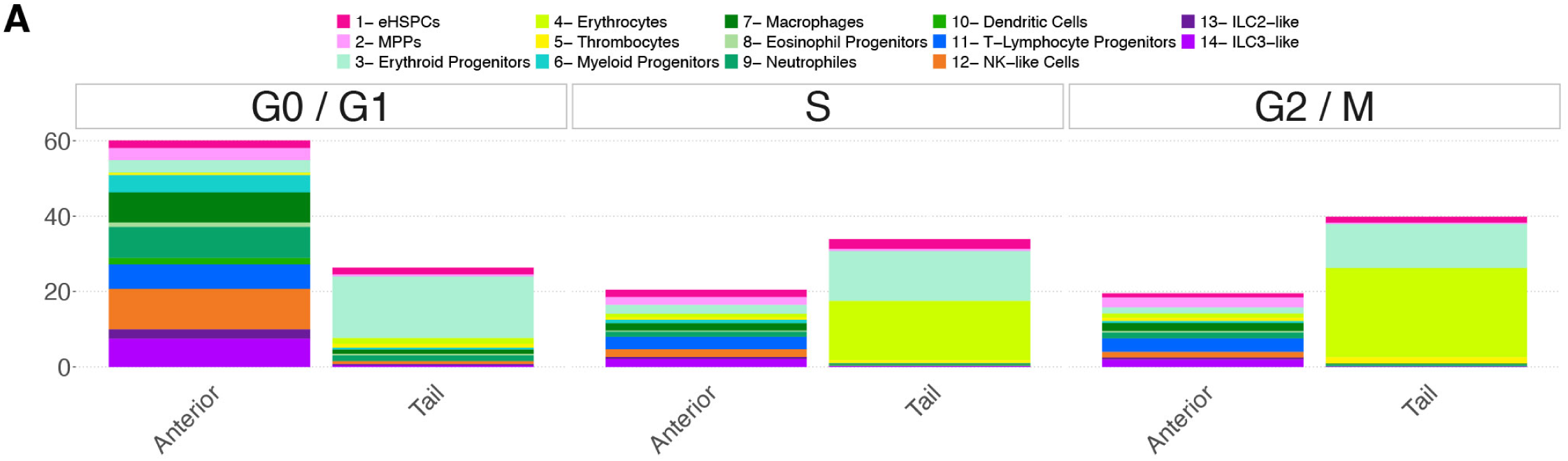
Cycling cells in the tail mostly belong to the erythroid lineage. Proportion of cell populations at different cell cycle stages in the anterior and posterior region of 5dpf larvae (Chromium dataset), mean values are represented.

**Supplementary Figure 5:**
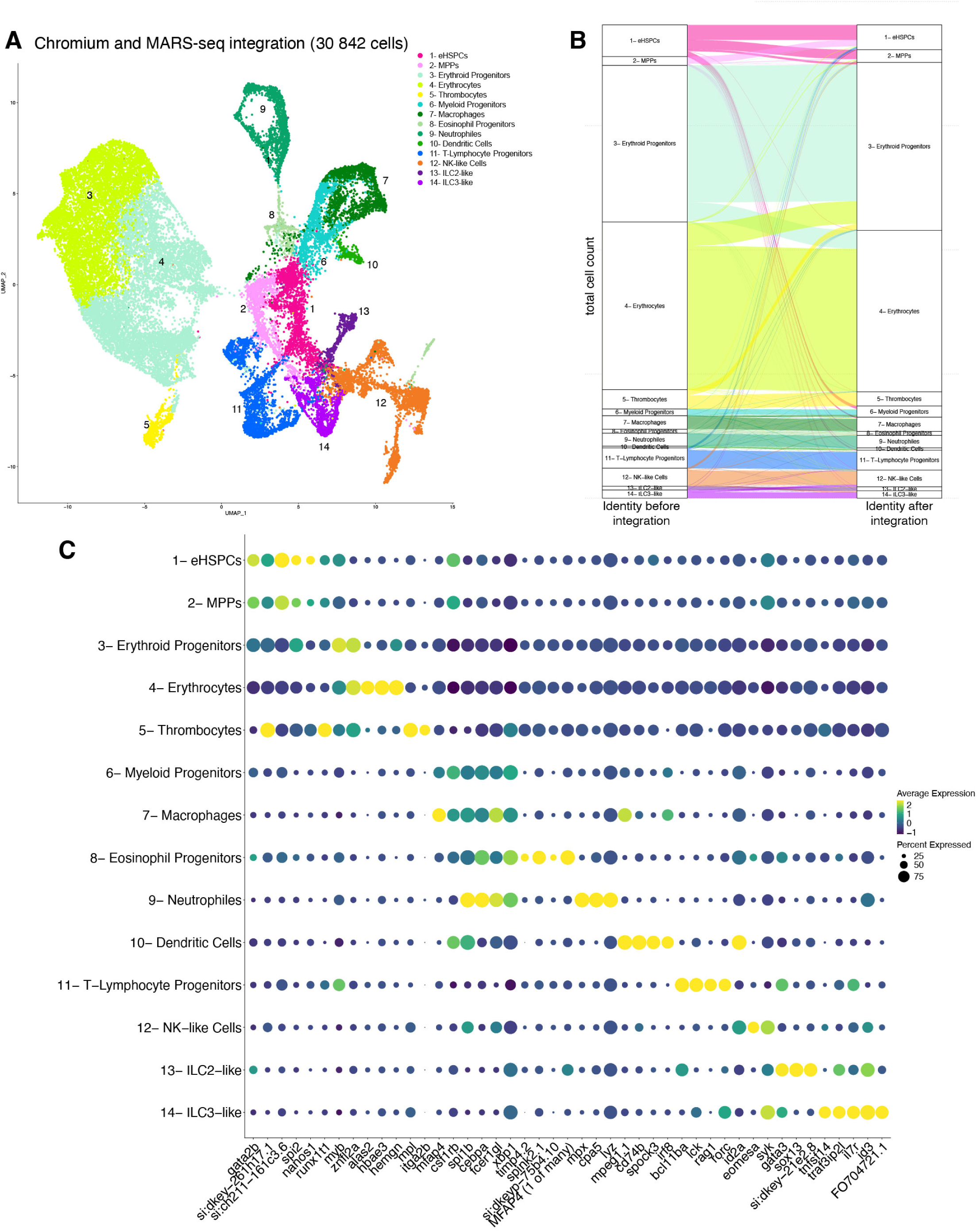
Integrated analysis of multiple technical and biological modalities for the in-depth characterization of hematopoietic larval populations. **(A)** UMAP of hematopoietic cells resulting from the integration of the Chromium and MARS-seq datasets. **(B)** Comparison of cell type assignation per cell before integration (Chromium and MARS-seq datasets, left) and within the integrated dataset (right). Overall, manual cell type assignation based on differential gene expression is globally reproduced after integration, with variability for closely related cell types (such as erythroid progenitors and erythrocytes or eHSPCs and MPPs). **(C)** Dot-plot of selected marker genes differentially expressed across defined hematopoietic populations.

**Supplementary Figure 6:**
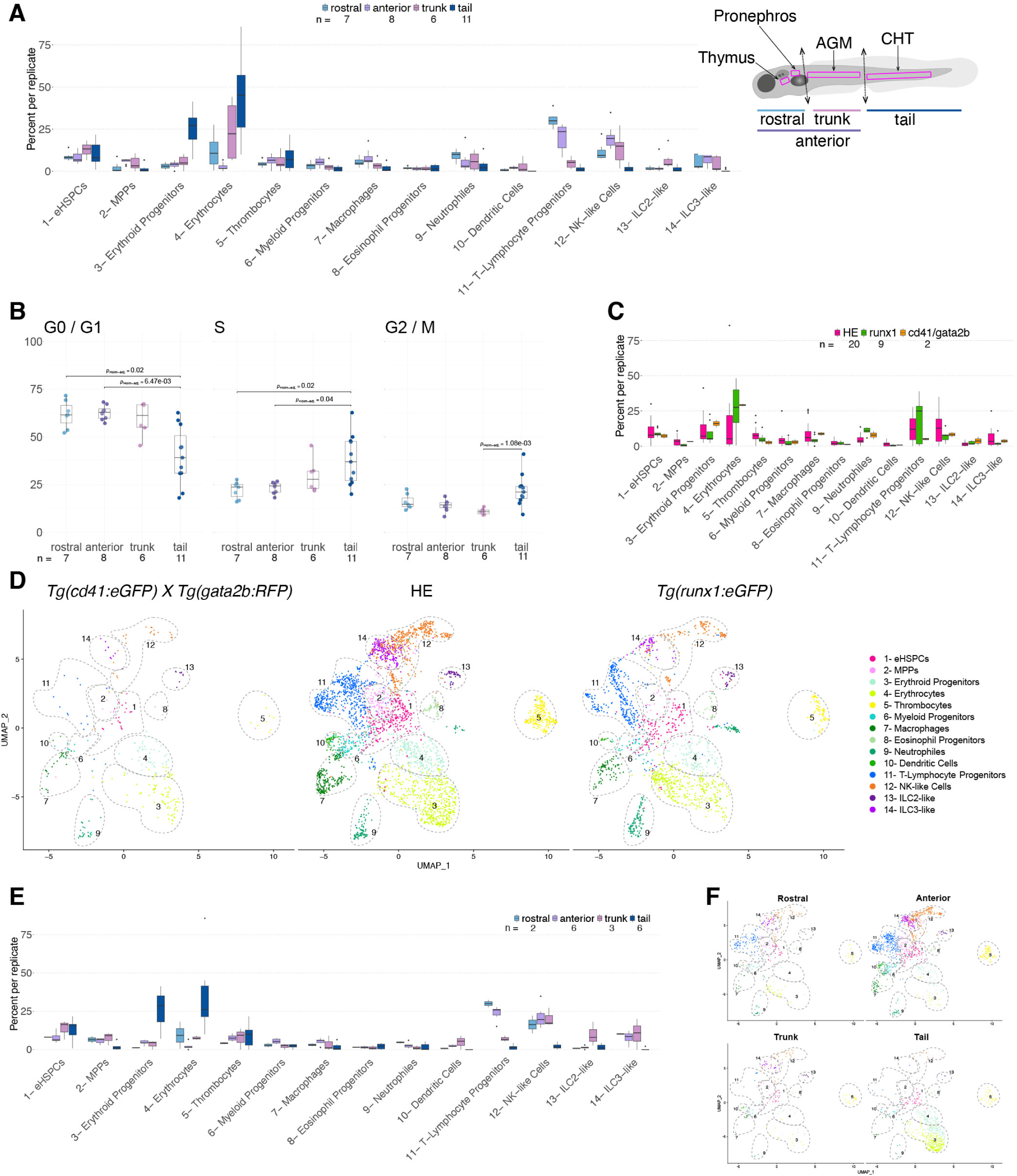
Comparative transcriptional analysis of hematopoietic populations across different transgenic origins and residing in different niches. **(A)** Proportion of hematopoietic populations within the different regions of the larva (right cartoon), in the total MARS-seq dataset. **(B)** Distribution of cells at each cell cycle stage within the different regions of the larva, in the MARS-seq dataset. **(C)** Proportion of hematopoietic populations for the different origins, in the MARS-seq dataset (see Fig. 4A). HE: photoconverted Hemogenic Endothelial (and EHT) progenies, *Tg(runx1+23:eGFP), Tg(cd41:eGFP; gata2b:RFP)*. **(D)** UMAPs of hematopoietic cells from the MARS-seq dataset, split by origin (same as in **C**). **(E)** Proportion of hematopoietic populations of the HE subset of the MARS-seq dataset within the different larval regions. **(F)** UMAPs of hematopoietic cells of the HE subset of the MARS-seq dataset, split by region. **(A, B)** n=7 replicates for rostral, n=8 for anterior, n=6 for trunk and n=11 for tail regions. **(B)** Holms adjusted two sided Wilcoxon tests, only significant p-value are shown**. (C, D)** n=20 replicates for HE, n=9 for runx1 and n=2 for *cd41/gata2b*. **(E, F)** n=2 replicates for rostral, n=6 for anterior, n=3 for trunk and n=6 for tail regions.

**Supplementary Figure 7:**
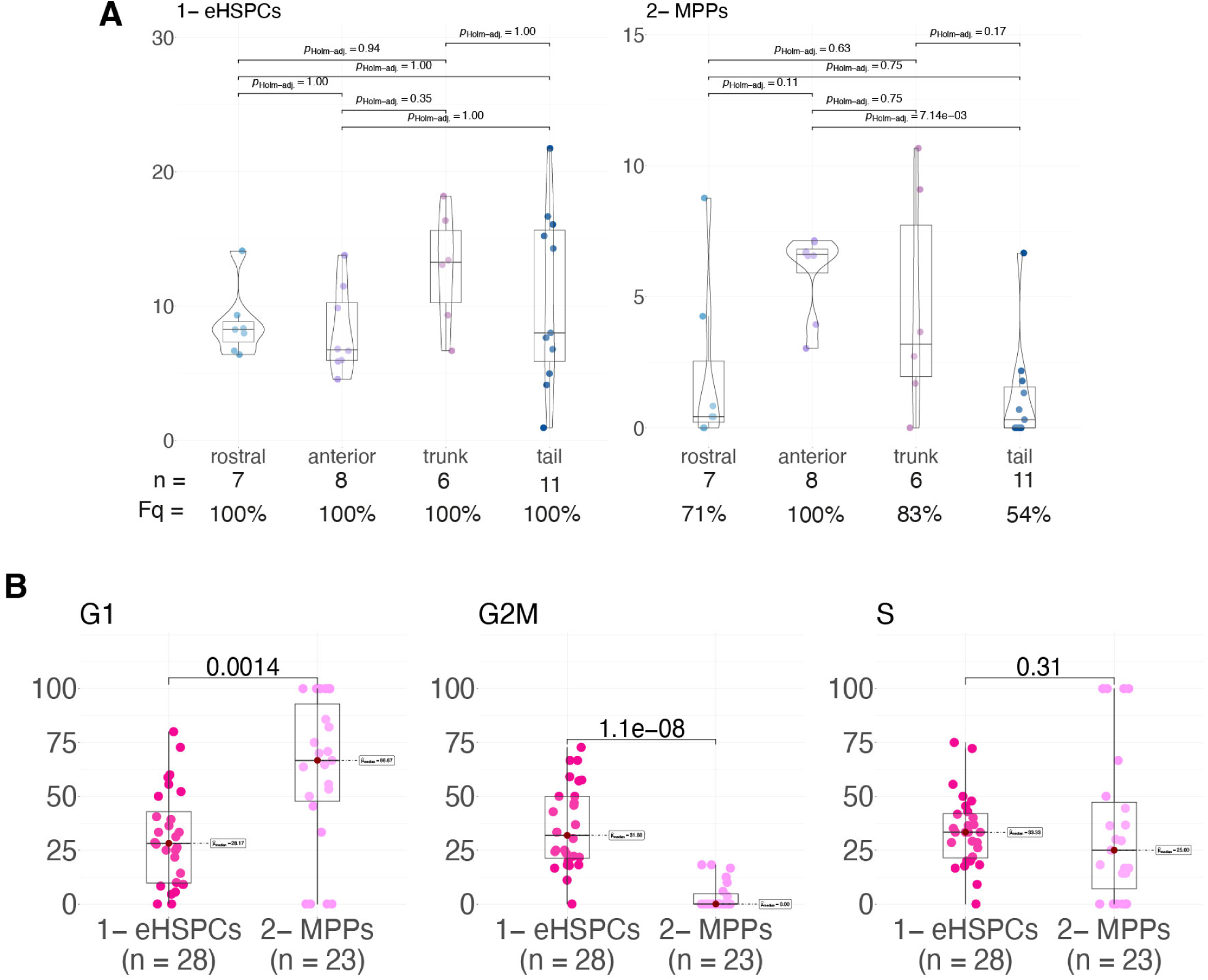
Cell cycle status of eHSPCs and MPPs and their repartition in developmental niches. **(A)** Percentage of eHSPC and MPP cells (MARS-seq dataset) in the different regions of the larva (rostral, trunk and tail, 7, 8, 6 and 11 replicates respectively). Fq: frequency of observation across replicates. Holms adjusted two sided Wilcoxon tests. **(B)** Percentage of eHSPC and MPP cells (MARS-seq dataset) in cell cycle phases (G0/G1, S and G2/M), n=28 and n=23 replicates for eHSPCs and MPPs respectively, two sided Wilcoxon tests.

**Supplementary Figure 8:**
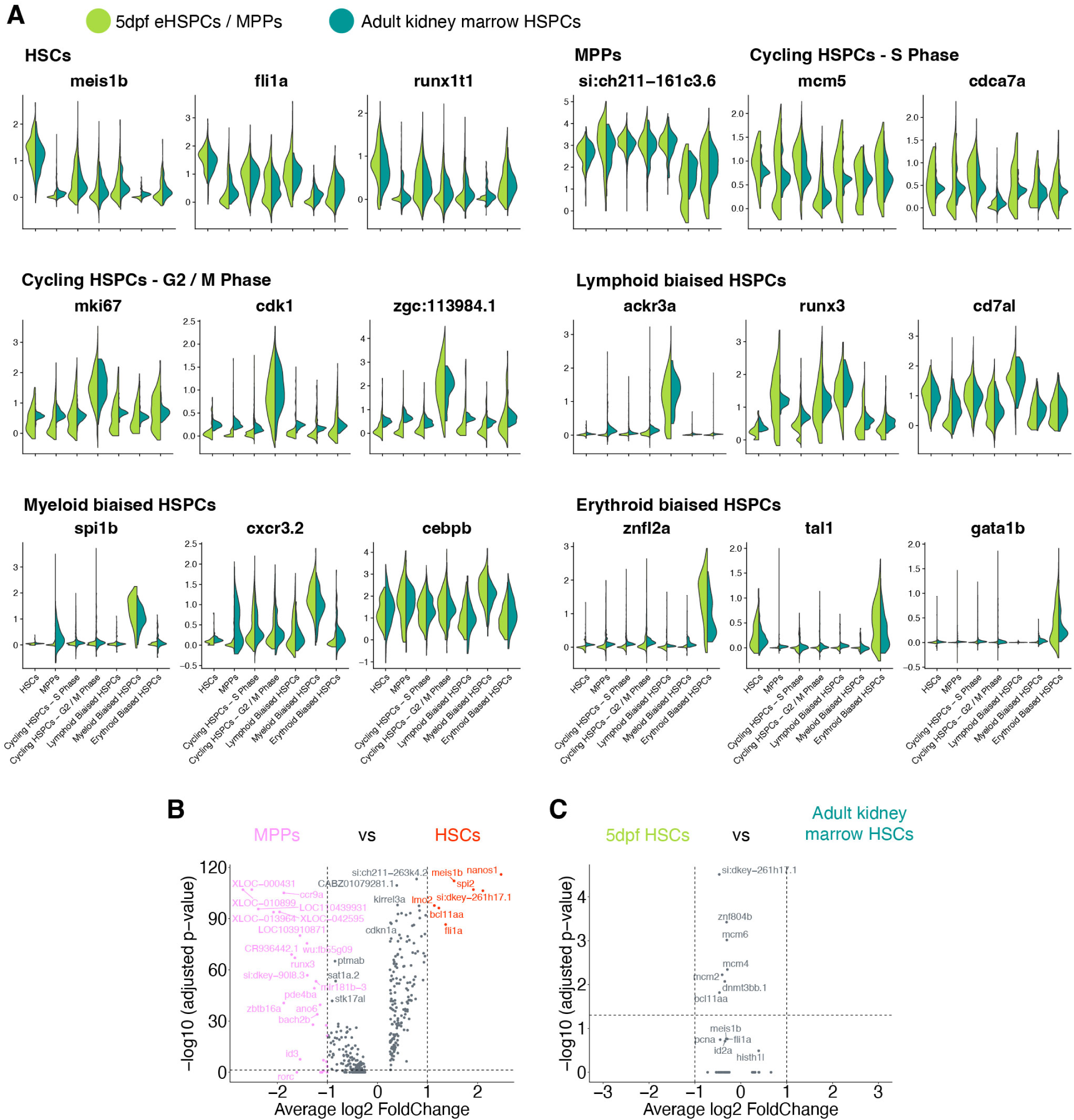
Characterization of HSPC populations found in developmental and adult datasets. **(A)** Violin plots of differentially expressed genes (non-parametric Wilcoxon rank sum test) specific of HSPC sub-populations characterized in Fig. 5D. **(B)** Differentially expressed genes between HSCs and MPPs of mixed developmental (5dpf whole larva) and adult (3 months kidney marrow) origins (Rubin et al., 2022). **(C)** Differentially expressed genes between HSCs of developmental (5dpf whole larva) and adult (3 months kidney marrow) origins. No genes are found to be differentially expressed between the two conditions, reinforcing their strong similarity in identity. **(B, C)** Non-parametric Wilcoxon rank sum test, threshold for p-value and log2 Fold change of 0.05 and 1 respectively.

**Supplementary Figure 9:**
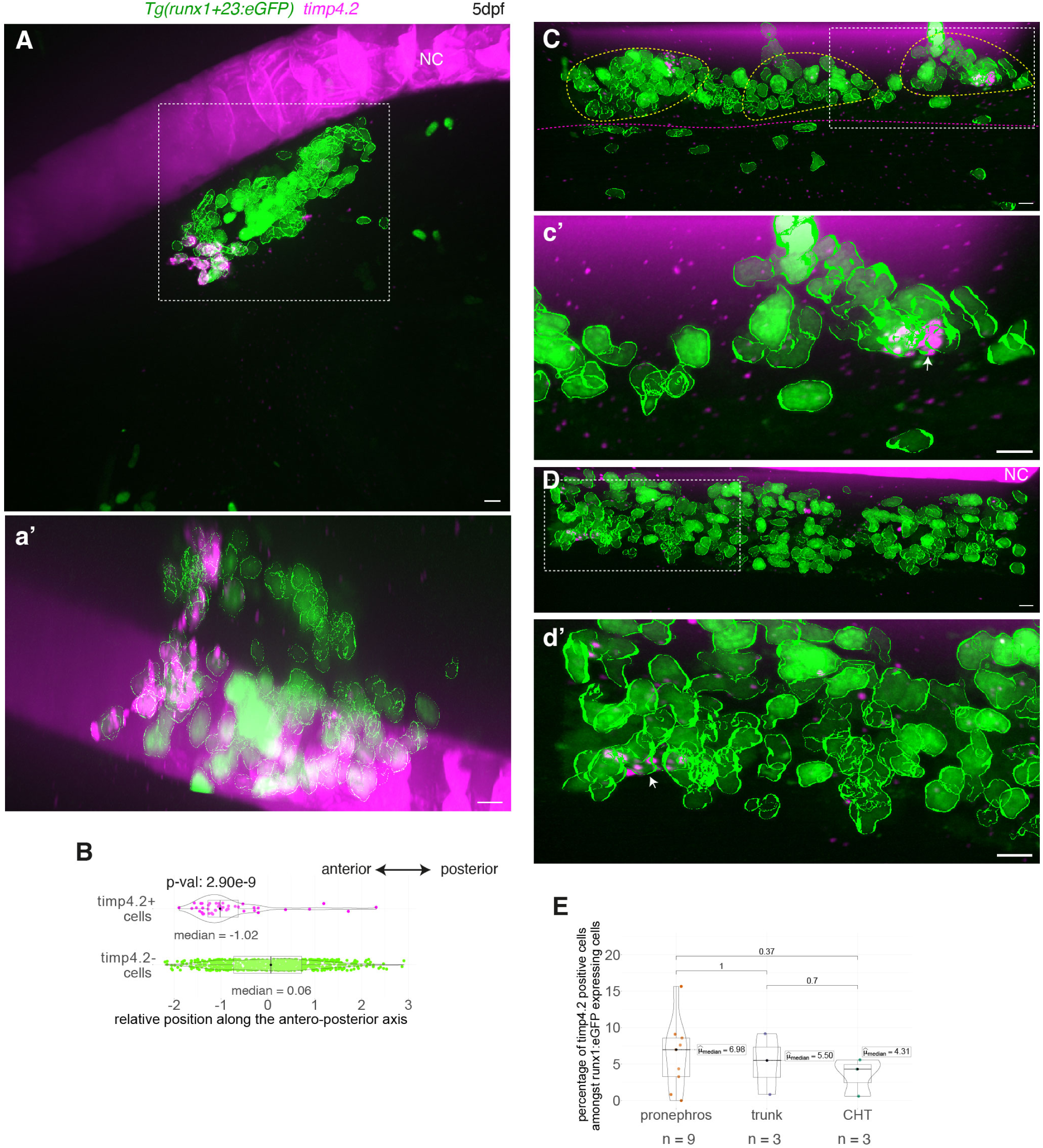
Whole mount *in situ* analysis of *timp4*.2 mRNA expression in hematopoietic and vascular cells using RNAscope. **(A, C, D)** Representative images (Imaris 3D-rendering) of RNAscope ISH for *timp4.2* (magenta spots) in 5dpf *Tg(runx1+23:eGFP)* larvae. Images show the pronephros region (**A**, see also **Movie 6**), the posterior trunk region (**C**, above the elongated yolk) and the CHT **(D)**. **(a’, c’, d’)** are magnifications of regions outlined with white dashed boxes in **(A, C, D),** respectively. eGFP+ hematopoietic cells were segmented (green contours). White arrows point at *timp4.2* positive hematopoietic cells. The sub-aortic clusters are delimited by yellow dashed lines and the gut by magenta dashed lines. **(B)** Relative position of eGFP+ cells along the antero-posterior axis of the pronephros (n=681 *timp4.2*-cells, n=46 *timp4.2*+ cells). **(E)** Percentage of eGFP+ hematopoietic cells expressing *timp4.2*, n=6 larvae for pronephros, n=3 for trunk and CHT regions. **(B, E)** Two sided Wilcoxon tests. NC: notochord. Scale bars: 10µm.

## Supplementary Movies

**Movie 1 (Relative to Fig. 2A):** 3D surface-rendering images of thymi and progenies of photoconverted EHT pol- (left) and EHT pol+ (right) cells observed in the thymus region of *Tg(kdrl:nls-kikume)* larvae, at 5dpf. 3 representative replicates are shown for each condition. The contour of the thymus has been segmented, based on green cell surface (outlined in grey). In magenta: progenies of photoconverted cells. Scale bars: 10µm.

**Movie 2 (Relative to Fig. 6A):** 3D visualization of RNAscope *in situ* hybridizations for *gata2b* (in magenta) in the pronephros region of *Tg(runx1+23:eGFP)* 5dpf larvae, 3 representative replicates are shown. Bottom row shows magnifications of the top row. Hematopoietic cells in the pronephros are delineated (green contours). Scale bars: 10µm.

**Movie 3 (Relative to Fig. 6D, E):** 3D visualization of RNAscope *in situ* hybridizations for *gata2b* (in magenta) in the lower trunk (top row) and CHT (bottom row) regions of *Tg(runx1+23:eGFP)* 5dpf larvae; representative images from 2 different larvae are shown. Hematopoietic cells are delineated (green contours). Scale bars: 10µm.

**Movie 4 (Relative to Fig. 6C):** 3D visualization of RNAscope *in situ* hybridizations for *gata2b* (in magenta) in the upper trunk region of *Tg(runx1+23:eGFP)* 5dpf larvae, representative images from 3 different larvae are shown. Hematopoietic cells are delineated (green contours). Scale bars: 10µm.

**Movie 5 (Relative to Fig. 6G, I):** 3D visualization of RNAscope *in situ* hybridizations (in magenta) for *gata2b* (top row) or *cmyb* (bottom row) in the upper trunk region of *Tg(kdrl:eGFP)* 5dpf larvae, representative images from 3 different larvae are shown. The contours of eGFP+ cells (endothelial cells and potentially newly generated hematopoietic cells) are shown, and color coded based on their localization (yellow for cells in and around the aorta, blue for cells of and in the vein plexus, white for cells in and around the SIA and green for unclassified cells). Scale bars: 10µm.

**Movie 6 (Relative to Fig. Supp9A):** 3D visualization of RNAscope *in situ* hybridizations for *timp4.2* (in magenta) in the pronephros region of *Tg(runx1+23:eGFP)* 5dpf larvae, 3 representative replicates are shown. Bottom row shows magnifications of the top row. Hematopoietic cells in the pronephros are delineated (green contours). Scale bars: 10µm.

**Movie 7 (Relative to Fig. 7A):** 3D visualization of RNAscope *in situ* hybridizations for *cd34* (in magenta) in the pronephros region of *Tg(runx1+23:eGFP)* 5dpf larvae, 3 representative replicates are shown. Bottom row shows magnifications of the top row. Hematopoietic cells in the pronephros are delineated (green contours). Scale bars: 10µm.

**Movie 8 (Relative to Fig. 7C, D):** 3D visualization of RNAscope *in situ* hybridizations for *cd34* (in magenta) in the trunk (top row) and CHT (bottom row) region of *Tg(runx1+23:eGFP)* 5dpf larvae, representative images from 2 different larvae are shown. Hematopoietic cells, with green contours, are shown. Scale bars: 10µm.

## Supplementary Tables

**Supplementary Table 1:** Differentially expressed genes per cluster and cell types for integrated Chromium dataset

**Supplementary Table 2:** Marker genes references for cell type identity assignation

**Supplementary Table 3:** Cell count per condition and replicate for MARS-seq analysis

**Supplementary Table 4:** Differentially expressed genes per cluster and cell types for integrated MARS-seq dataset

**Supplementary Table 5:** Differentially expressed genes per cluster and cell types for integrated Chromium and MARS-seq datasets

**Supplementary Table 6:** Differentially expressed genes per cluster and cell types for integrated Chromium and Adult HSPCs (Rubin et al., 2022) datasets. Comparison of gene expression between integrated HSCs and MPPs clusters. Comparison of gene expression between developmental and adult HSCs.

## Author contributions

**LT:** Conceptualization, Methodology, Software, Validation, Formal analysis, Investigation, Resources, Data curation, Writing – original draft preparation, Writing – review and editing, Visualization; **CV:** Methodology, Investigation; **SS:** Methodology, Validation, Investigation, Resources; **YL-M:** Methodology, Software, Formal analysis, Investigation; **AS:** Conceptualization, Methodology, Software, Validation, Formal analysis, Investigation, Resources, Data Curation, Writing – original draft preparation, Writing – review and editing, Visualization, Supervision, Project administration, Funding acquisition

## Acknowledgements

We wish to warmly thank Philippe Herbomel allowing the initiation of this project and providing resources for the photoconversion station (grant from the Fondation pour la Recherche Médicale (#DEQ20160334881)). We thank our fish facility members Yohann Rolin and Karim Sebastien for their daily help and commitment. We thank Laure Bally-Cuif for her support. We wish to thank David Traver for providing the *gata2b* transgenic fish line. We thank Olivier Mirabeau, Baptiste Saudemont, Marc Monot and David Morizet for their expertise and invaluable assistance in single-cell analyses and bio-informatics. We acknowledge the help of the Cytometry and Biomarkers (UTechS CB) service platform for expert assistance with FACS analysis and sc-RNA-seq technologies and access to equipment. We acknowledge the help of the Image Analysis Hub, of the Photonic BioImaging (UTechS PBI), of the Biomics and of the Bioinformatics and Biostatistics Hub service platforms of the Institut Pasteur for their respective contributions.

## Fundings

This work was supported by Institut Pasteur, CNRS, grants from La Ligue Contre le Cancer, Comité de Paris (RS21/75-5 and RS22/75-9) to AS, grant from the CNRS GDR3740 “Stem cells *in vivo*” to LT. LT was recipient of a PhD fellowships from the Collège Doctoral, Sorbonne Université, and from the Labex Revive (ANR-10-LABX-73).

## Competing interests

The authors declare no competing or financial interests.

## Data availability statement

The following datasets have been generated for this work. They have been deposited on the open external repository Zenodo (https://zenodo.org/). Access to the source data on the repository is restricted until acceptation of the manuscript.

**Source data 1** relative to Fig. 2 and **Movie 1,** confocal z-stack acquisitions, single-cell tracing (photoconversion), https://doi.org/10.5281/zenodo.13886928

**Source data 2** relative to **Figs 3-5**, **S2-S8,** newly generated sc-RNA-seq datasets (MARS-seq, Chromium and integrated MARS-seq/Chromium dataset), source data table, figure plotting scripts. https://doi.org/10.5281/zenodo.13904330

**Source data 3** relative to Fig. 6A and **Movie 2,** confocal z-stack acquisitions of RNAscope *in situ* hybridization (*gata2b*), https://doi.org/10.5281/zenodo.13884530

**Source data 4** relative to Fig. 6A and **Movie 2,** confocal z-stack acquisitions of RNAscope *in situ* hybridization (*gata2b*), https://doi.org/10.5281/zenodo.13884643

**Source data 5** relative to Fig. 6D, E and Movie 3, confocal z-stack acquisitions of RNAscope *in situ* hybridization (*gata2b*), https://doi.org/10.5281/zenodo.13884760

**Source data 6** relative Fig. 6C, G and **Movie 4, 5,** confocal z-stack acquisitions of RNAscope *in situ* hybridization (*gata2b*), https://doi.org/10.5281/zenodo.13884837

**Source data 7** relative to Fig. 6I and **Movie 5**, confocal z-stack acquisitions of RNAscope *in situ* hybridization (*cmyb*), https://doi.org/10.5281/zenodo.13884901

**Source data 8** relative to **Fig. S9A** and **Movie 6,** confocal z-stack acquisitions of RNAscope *in situ* hybridization (*timp4.2*), https://doi.org/10.5281/zenodo.13884936

**Source data 9** relative Fig. 7A, C, D and **Movie 7, 8,** confocal z-stack acquisitions of RNAscope *in situ* hybridization (*cd34*), https://doi.org/10.5281/zenodo.13885925

## Notes

### Competing Interest Statement

The authors have declared no competing interest.

